# Muscle weakness and mitochondrial stress occur before metastasis in a novel mouse model of ovarian cancer cachexia

**DOI:** 10.1101/2024.04.08.588639

**Authors:** Luca J. Delfinis, Leslie M. Ogilvie, Shahrzad Khajehzadehshoushtar, Shivam Gandhi, Madison C. Garibotti, Arshdeep K. Thuhan, Kathy Matuszewska, Madison Pereira, Ronald G. Jones, Arthur J. Cheng, Thomas J. Hawke, Nicholas P. Greene, Kevin A. Murach, Jeremy A. Simpson, Jim Petrik, Christopher G.R. Perry

## Abstract

**Objectives:** A high proportion of women with advanced epithelial ovarian cancer (EOC) experience weakness and cachexia. This relationship is associated with increased morbidity and mortality. EOC is the most lethal gynecological cancer, yet no preclinical cachexia model has demonstrated the combined hallmark features of metastasis, ascites development, muscle loss and weakness in adult immunocompetent mice.

**Methods:** Here, we evaluated a new model of ovarian cancer-induced cachexia with the advantages of inducing cancer in adult immunocompetent C57BL/6J mice through orthotopic injections of EOC cells in the ovarian bursa. We characterized the development of metastasis, ascites, muscle atrophy, muscle weakness, markers of inflammation, and mitochondrial stress in the tibialis anterior (TA) and diaphragm ∼45, ∼75 and ∼90 days after EOC injection.

**Results:** Primary ovarian tumour sizes were progressively larger at each time point while robust metastasis, ascites development, and reductions in body, fat and muscle weights occurred by 90 Days. There were no changes in certain inflammatory (TNFα), atrogene (MURF1 and Atrogin) or GDF15 markers within both muscles whereas IL-6 was increased at 45 and 90 Day groups in the diaphragm. TA weakness in 45 Day preceded atrophy and metastasis that were observed later (75 and 90 Day, respectively). The diaphragm demonstrated both weakness and atrophy in 45 Day. In both muscles, this pre-metastatic muscle weakness corresponded with considerable reprogramming of gene pathways related to mitochondrial bioenergetics as well as reduced functional measures of mitochondrial pyruvate oxidation and creatine-dependent ADP/ATP cycling as well as increased reactive oxygen species emission (hydrogen peroxide). Remarkably, muscle force per unit mass at 90 days was partially restored in the TA despite the presence of atrophy and metastasis. In contrast, the diaphragm demonstrated progressive weakness. At this advanced stage, mitochondrial pyruvate oxidation in both muscles exceeded control mice suggesting an apparent metabolic super-compensation corresponding with restored indices of creatine-dependent adenylate cycling.

**Conclusion:** This mouse model demonstrates the concurrent development of cachexia and metastasis that occurs in women with EOC. The model provides physiologically relevant advantages of inducing tumour development within the ovarian bursa in immunocompetent adult mice. Moreover, the model reveals that muscle weakness in both TA and diaphragm precedes metastasis while weakness also precedes atrophy in the TA. An underlying mitochondrial bioenergetic stress corresponded with this early weakness. Collectively, these discoveries can direct new research towards the development of therapies that target pre-atrophy and pre-metastatic weakness during EOC in addition to therapies targeting cachexia.

**Highlights:**

- This study reports the first orthotopic model of metastatic ovarian cancer cachexia that can be induced in adult immunocompetent mice
- Diaphragm and limb muscle weakness precedes metastasis and atrophy during ovarian cancer
- Skeletal muscle mitochondrial oxidative and redox stress signatures occur during pre-metastatic stages of ovarian cancer
- Specific muscle force as well as mitochondrial pyruvate oxidation and creatine metabolism demonstrate compensation in later stages
- Ovarian cancer has heterogeneous effects on distinct muscle types across time

## 1. Introduction

Cancer-induced cachexia is a multifactorial syndrome characterized by muscle loss and weakness [1]. Severe cachexia is linked to reductions in quality of life, tolerance to anticancer therapies and overall survivability [2–4]. The prevalence of cachexia varies widely (20-80%) across different types and severity of cancer [5,6] raising the possibility that cachexia may have both ubiquitous and distinct mechanisms related to the host organ. With growing awareness that cancer itself can induce cachexia even in the absence of cytotoxic cancer therapies, it is imperative to develop pre-clinical models for each type of cancer cachexia that captures critical phenotypic hallmarks of this disease in humans.

To date, there are limited pre-clinical models available for ovarian cancer-induced cachexia despite epithelial ovarian cancer (EOC) being the most lethal gynecological cancer in women [7]. To our knowledge, only three preclinical studies have evaluated ovarian cancer-induced cachexia using a combination of *in vitro* and animal models [8–10] with the latter involving either a genetic mutation, whereby mice are born with ovarian cancer or injections of human-derived ovarian cancer cells under the skin of immunodeficient mice. While both genetic and subcutaneous injection approaches are generally used to research many types of cancer in animal models [11,12] and have provided significant advancements in our understanding of cancer-induced cachexia, there remains a need to develop a model enabling tumour-induction into the host ovaries at selected ages in adult mice. The development of a new preclinical model would ideally capture other critical features of cancer cachexia including metastasis given this defining event is associated with severe cachexia and reduced survival rates during advanced stages of ovarian cancer [13]. Indeed, when ovarian cancer is detected at early stages and before metastasis, the cure rate is estimated to be as high as 90% [14] in contrast to much lower survival rates once metastasis has occurred. As more than 70% of ovarian cancer cases are diagnosed at late stages, improving our understanding of how cachexia develops could lead to new insight into improved patient management and perhaps early detection of ovarian cancer itself [13]. Furthermore, metastasis is not always capitulated in a variety of cancer-specific cachexia preclinical models which presents a collective limitation for many areas of research [15]. In this regard, it has been suggested that metastatic, immunocompetent models with cancer cells injected into the host organ (orthotopic) designed for each type of cancer could greatly improve the predictive power of mouse models for both mechanistic investigations and pre-clinical therapy development [15].

Characterizing cachexia warrants careful consideration of how changes in muscle mass and force production occur over time as the tumour develops, and in relation to underlying mechanisms regulating both aspects of muscle quality. Recently, we reported that muscle weakness precedes atrophy in the C26 (Colon-26) colorectal mouse [16]. This pre-atrophy weakness also occurred without any changes in expression of classic atrophy-related gene programs suggesting muscle weakness during cancer could also be caused by unknown atrophy-independent mechanisms. In this study, a strong relationship was found between pre-atrophy weakness and mitochondrial pathway-specific reprogramming as an apparent early metabolic stress response to the initial appearance of tumours. Remarkably, once locomotor muscle atrophy occurred, mitochondria appeared to adapt by increasing pyruvate oxidation, which was related to an unexpected restoration of mass-specific force production [16]. Of interest, this relationship was heterogeneous across different types of muscles suggesting the effects of cancer on one muscle type do not necessarily predict the response in another muscle. This phenomenon demonstrates the value of comparing muscle force to muscle mass across time and between muscle types in relation to tumour size. In this regard, muscle weakness during the pre-atrophy and atrophy (cachexia) phases of ovarian cancer have not been investigated, nor in relation to metastasis or metabolic dysfunction. Understanding this could lead to more precise understanding of ovarian cancer cachexia pathology to aid better mechanism elucidation and therapy development.

The purpose of this study was to identify the time-dependent and muscle-specific development of weakness and atrophy in a novel model of ovarian cancer cachexia in relation to metabolic reprogramming. In this model, spontaneously transformed ovarian epithelial cells from the same mouse strain (syngeneic) were injected into the ovarian bursa (orthotopic) in immunocompetent mice with the intention of retaining the normal immune response to this type of cancer. These results demonstrate a new cachexia mouse model that captures metastasis characteristic of advanced stages of EOC in women [17,18] that also has considerable utility for identifying new relationships between the development of muscle weakness, atrophy, and metabolic reprogramming across time during ovarian cancer.

## 2. Materials and Methods

### 2.1 Animal Care and ID8 Inoculation

Two cohorts of 48 (n=12 per group; **SFigure 1A and 1B**) female C57BL/6 mice were ordered at 7-9 weeks of age from Charles River Laboratories. These mice were housed at the University of Guelph in accordance with the Canadian Council on Animal Care. Tumours were induced as described previously at the University of Guelph [19–22]. Briefly, ID8 cells (transformed murine epithelial cells; 1.0 x 10^6^ in 5μL) were injected directly under the left ovarian bursa of C57BL/6 mice generating an orthotopic, syngeneic, immunocompetent cancer mouse model. Control mice were sham injected with equivalent volumes of sterile phosphate buffered saline (PBS). Two weeks after ID8 inoculation, mice were transported from University of Guelph to York University where they were housed for the remainder of the study in accordance with the Canadian Council on Animal Care. All mice were provided access to standard chow and water *ad libitum*.

**Figure 1.**
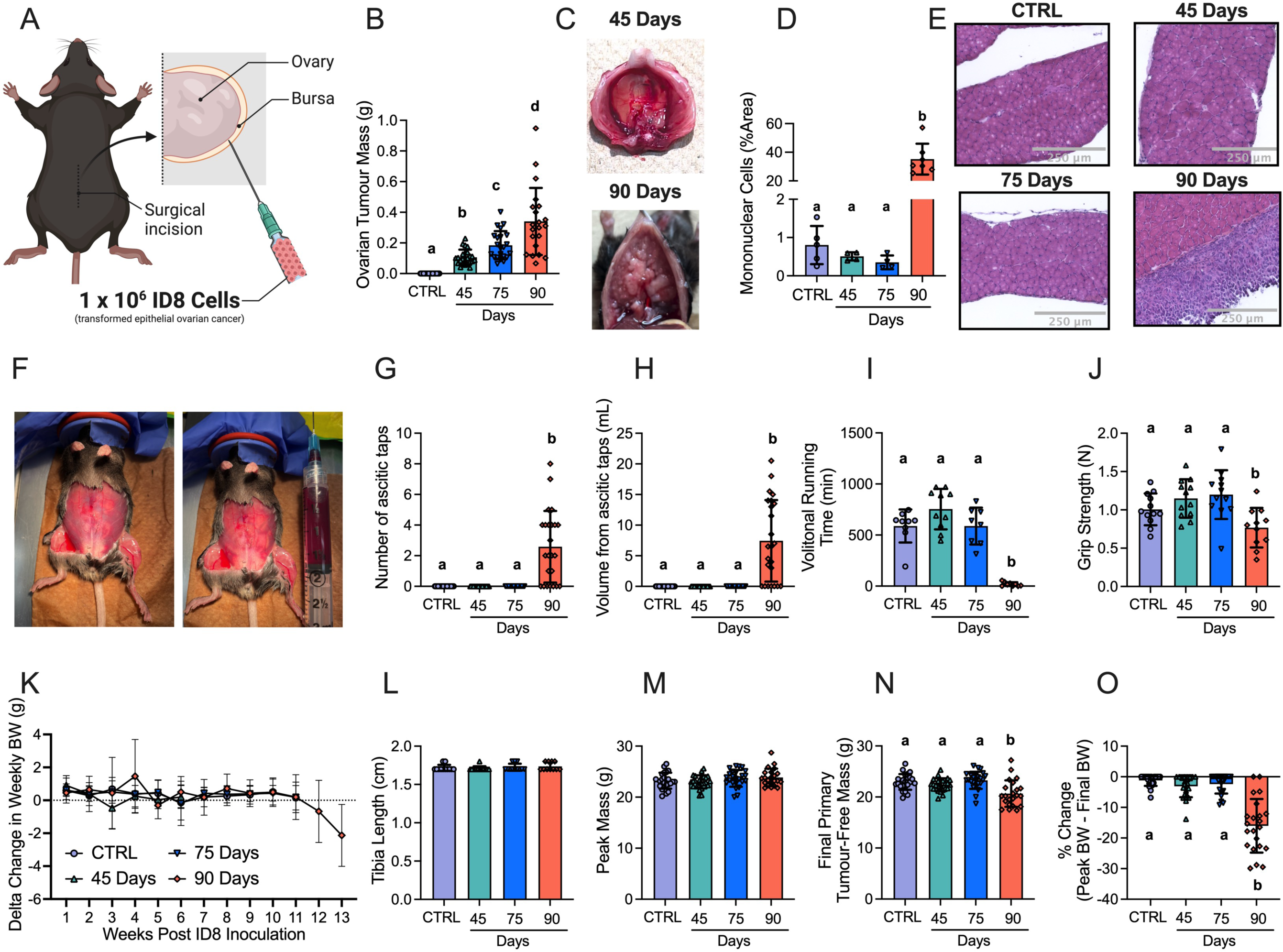
The effects of transformed epithelial ovarian cancer cells (ID8) implantation underneath the ovarian bursa of C57BL6 mice. 1 x 10^6^ ID8 cells were injected underneath the ovarian bursa (A) and developed for 42-48, 72-78 and 83-107 days (45 Day, 75 Day and 90 Day time points respectively). Control mice were injected with identical volumes of PBS and aged for 72-78 days. Primary ovarian tumour mass was measured at sacrifice (B, n = 21-24). Noticeable metastasis of ovarian cancer cells occurred by the 90-day time point and were photographed for qualitative assessment (C). Hematoxylin & eosin staining was used to assess mononuclear cell infiltration as an index of metastasis (D, n = 4-7, E Representative images; original magnification, x20). Mice developed ascites after ∼75 days of ovarian cancer (F) and were tapped to prolong their survival (G & H, n = 24). Volitional wheel running (I, n = 8-11) and grip strength (J, n = 11-12) were used to assess voluntary motor function. Body weights were also measured every week and the delta weekly body weight (BW) was analyzed (K, n = 22-24). Tibia length (L, n = 11-12), peak body weight (M, n = 22-24), and final primary ovarian tumour-free body weight (N, n = 22-24) were also assessed. Percent change from peak body weight to final body weight was analyzed (O, n = 22-24). Results represent mean ± SD. Lettering denotes statistical significance when different from each other (*p* < 0.05). All data was analyzed using a one-way ANOVA and followed by a two-stage step-up method of Benjamini, Krieger and Yukutieli multiple comparisons test. Data that was not normally distributed was analyzed with a Kruskal-Wallis test followed by the same post-hoc analysis. C57BL/6J female mice ∼75 days post PBS injection as controls (CTRL); C57BL/6J female mice ∼45 days post ovarian cancer injection (45 Days); C57BL/6J female mice ∼75 days post ovarian cancer injection (75 Days); C57BL/6J female mice ∼90 days post ovarian cancer injection (90 Days).

Control (CTRL) and 75 Day mice were injected at 9-11weeks old with PBS and ID8 cells respectively and aged for 72–78 days post injection. 45 Day mice were injected at 16-17 weeks old and aged for 42-48 days post injection. 90 Day mice were injected at 9-10 weeks old and aged for 83-107 days post injection (**SFigure 1A and 1B**). These ranges were chosen given force and mitochondrial assessments limit daily experimental throughput, and health metrics used to determine the date of euthanasia were variable in the more advanced stages of cancer. Specifically, at the 90 Day time point, mice were euthanized upon presentation of some of the following endpoint criteria: >10% body weight loss, >20mL of ascitic volume collected during paracentesis, > 5 ascites paracentesis taps completed, and/or subjective changes in behavioural patterns consistent with removal criteria as per animal care guidelines (self-isolation, ruffled fur/poor self-grooming and irregular gait). Staggering the age at which mice received cancer cells permitted a consistent age at euthanasia for all mice (20-24 weeks old) to reduce aging effects on all measures.

### 2.2 Volitional Wheel Running & Forelimb Grip Strength

72 hours before euthanasia, a subset of mice were placed in individual cages with a 14 cm diameter running wheel and rotation counter (VDO m3 bike computer, Mountain Equipment Co-Op, Vancouver, Canada) as done previously [23]. 24 hours later, distance and time ran were recorded and mice were placed in separate caging with no running wheel. Muscle measurements were made 48 hours thereafter. On the day of euthanasia, mice were removed from cages and brought towards a metal grid until such time the mice grasped the grid with the forepaws. Upon grasping, animals were pulled away from the grid until the grasp was released. Peak tension was recorded and this was repeated twice more with the maximum peak tension of 3 trials was used for analyses as done previously [23].

### 2.3 Tissue Collection Procedure

Mice were anesthetized with isoflurane and hearts were removed for euthanasia. All hindlimb muscles, inguinal subcutaneous fat and spleens were weighed and snap-frozen in liquid nitrogen and stored at −80^ο^C. Primary ovarian tumours were also collected by removing the ovary and tumour at the site of injection and carefully separating the tumour mass from the ovary mass and stored in liquid nitrogen. Hindlimb muscles were also embedded in optimal cutting temperature (OCT) medium and frozen (see section below). Tibialis anterior (TA) and diaphragm muscle were placed in BIOPS containing (in mM) 50 MES Hydrate, 7.23 K_2_EGTA, 2.77 CaK_2_EGTA, 20 imidazole, 0.5 dithiothreitol, 20 taurine, 5.77 ATP, 15 PCr, and 6.56 MgCl_2_·6 H_2_O (pH 7.1) to be prepared for mitochondrial bioenergetic assays.

### 2.4 Sectioning, histochemical & immunofluorescent staining

Tibialis anterior and diaphragm muscle samples were embedded in OCT medium (Thermo Fisher Scientific) and frozen in 2-methylbutane. These muscles were then sectioned into 10μm sections with a cryostat (HM525 NX, Thermo Fisher Scientific) maintained at −20^ο^C on Fisherbrand Superfrost Plus slides (Thermo Fisher Scientific). Hematoxylin and eosin (H&E) staining was used to assess mononuclear cell infiltration. Images were taken using EVOS M7000 imager (Thermo Fisher Scientific) using 20x magnification and analyzed on ImageJ. Immunofluorescent analysis of myosin heavy chain (MHC) expression was completed as previously described [24]. Images were taken with EVOS M7000 equipped with standard red, green and blue filter cubes. Fibers that did not fluoresce were considered IIx fibers. A total of 25-50 fibers were then randomly selected and measured. Type IIb fibers in the diaphragm strips were in low abundance, therefore, 5-27 fibers were measured and used for analysis. Type I fibers were extremely low in abundance in the TA and thus were not analyzed [24]. These images were also analyzed for cross sectional area (CSA) on ImageJ in a blinded fashion. Immunofluorescent analysis of embryonic myosin heavy chain (eMHC) were adapted from previous literature [25]. Briefly, sections were fixed with 10% formalin, blocked with 10% goat serum, followed by mouse IgG block (BML 2202; Vector Laboratories Inc., Burlingame, CA), and incubated with anti-eMHC (15μg/mL; DHSB F1.652) overnight. Secondary Alexa Fluor 647 IgG (1:1000; Abcam, ab150107) was then used to fluoresce eMHC primary antibody. Sections were then re-blocked once again and incubated with wheat-germ agglutinin (WGA; 1:1000; Invitrogen W11261) pre-conjugated to Alexa Fluor 488. Last, samples were mounted with DAPI mounting medium (Abcam, ab104139). D2.*mdx* muscle tissue saved from previous studies in our lab was used to evaluate the efficacy of the antibodies (**SFigure 2**).

**Figure 2.**
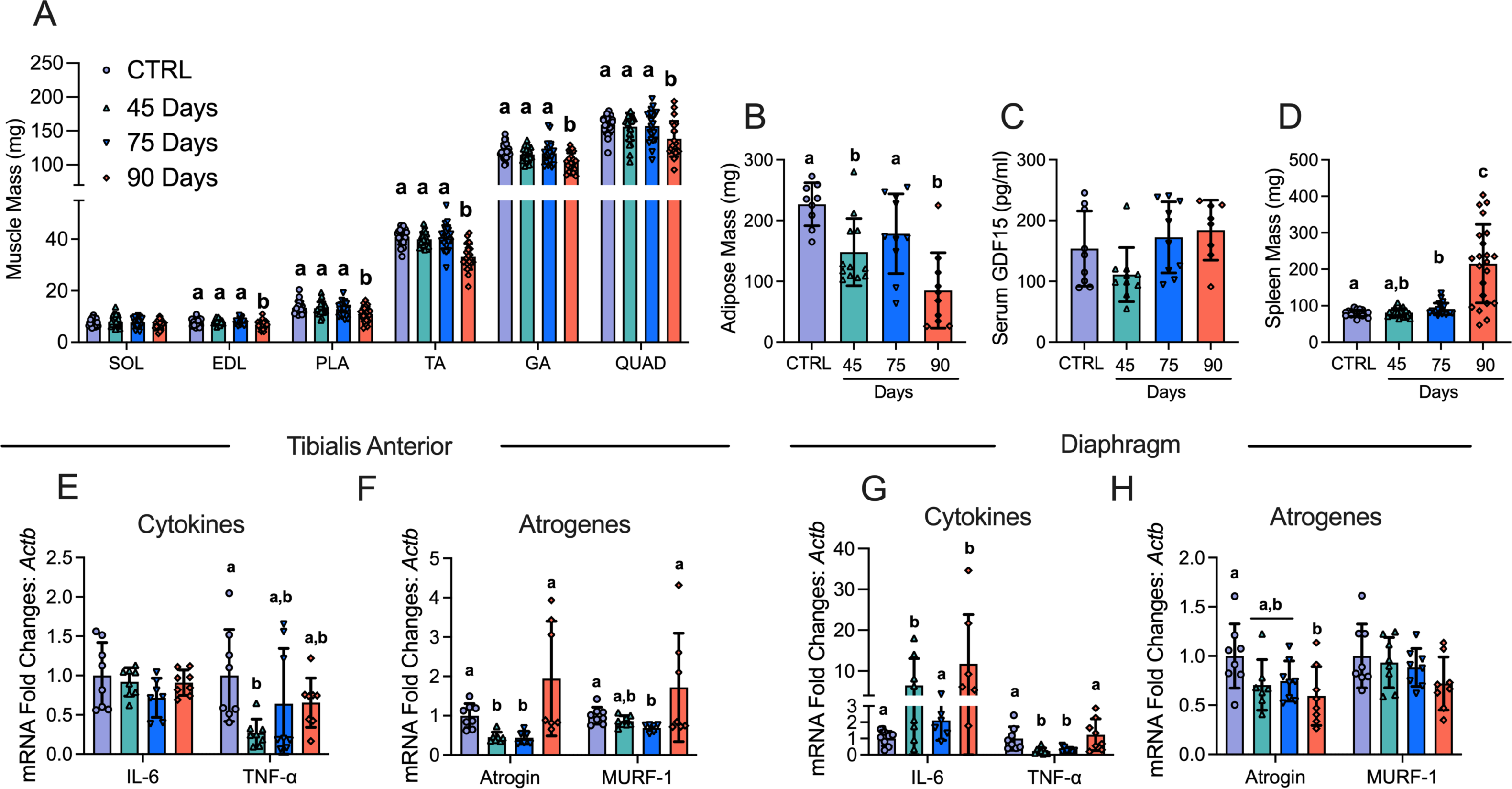
The effects of ID8 implantation on muscle mass, fat mass, spleen mass, GDF15 and gene expression of inflammation and atrogenes. Analysis of muscle mass at all time points in hindlimb muscles was completed (A, n = 22-24; soleus (SOL), extensor digitorum longus (EDL), plantaris (PLA), tibialis anterior (TA), gastrocnemius (GA) and quadriceps (QUAD)). Subcutaneous adipose mass in the inguinal fat depot (B, n = 9-12), serum GDF15 (C, n=8-11) and spleen mass (D, n = 21-22) were also analyzed. mRNA content of inflammatory and atrophy markers interleukin-6 (IL-6), tumour necrosis factor – alpha (TNF-α), atrogin and muscle RING-finger protein-1 (MURF-1) were measured using quantitative PCR in the TA and diaphragm of all groups (E-H, n = 6-8). Results represent mean ± SD. Lettering denotes statistical significance when different from each other (*p* < 0.05). C57BL/6J female mice ∼75 days post PBS injection as controls (CTRL); C57BL/6J female mice ∼45 days post ovarian cancer injection (45 Days); C57BL/6J female mice ∼75 days post ovarian cancer injection (75 Days); C57BL/6J female mice ∼90 days post ovarian cancer injection (90 Days). All data was analyzed using a one-way ANOVA or Kruskal-Wallis test when data did not fit normality. All ANOVAs were followed by a two-stage step-up method of Benjamini, Krieger and Yukutieli multiple comparisons test.

### 2.5 In Situ Tibialis Anterior Force and In Vitro Diaphragm Force

*In situ* TA force production was partially adapted from previous literature [26,27]. Mice were anesthetized with isoflurane and the distal tendon of the TA was exposed by incision at the ankle. The distal tendon was tied with suture thread as close to the muscle attachment as possible. Once the knot was secured the distal tendon was severed. Small cuts were made up the lateral side of the TA to expose the muscle for needle electrode placement. The knot was tied to an Aurora Scientific 305C (Aurora Scientific Inc., Aurora, ON, Canada) muscle lever arm with a hook. The foot of the mouse was secured with tape and the knee was immobilized with a needle and set screw with the length of the limb parallel to the direction of force. The two needle electrodes were placed in the gap of fascia between the TA and tibia to stimulate the common peroneal nerve (10-50 mA). The mouse was heated with a heating pad or heat lamp throughout force collection. Optimal resting length (L_o_) was determined using single twitches (pulse width=0.2ms) at 1 Hz stimulation frequency with 1 minute rest in between contractions to avoid fatigue. Once L_o_ was established, a ruler was used to determine length before the start of force-frequency collection (1, 10, 20, 30, 40, 50, 60, 80, 100, 120 and 200Hz with 1 minute rest in between contractions). Force production was normalized to the calculated CSA of the muscle strip (m/l*d) where m is the muscle mass, l is the length, and d is mammalian skeletal muscle density (1.06mg.mm^3^).

*In vitro* force production for diaphragm muscle was done as completed previously [16,28,29]. Briefly, the diaphragm strip was carefully sutured in Ringer’s solution (containing in mM: 121 NaCl, 5 KCl, 1.8 CaCl_2_, 0.5 MgCl_2_ 0.4 NaHPO_4_, 24 NaHCO_3_, 5.5 glucose and 0.1 EDTA; pH 7.3 oxygenated with 95% O_2_ and 5% CO_2_) such that the thread secured to the central tendon and ribs. The strip was then placed in an oxygenated bath filled with Ringer’s and maintained at 25^ο^C. The suture secured to the central tendon was then attached to the lever arm and the loop secured to the ribs was attached to the force transducer. The strip was situated between flanking platinum electrodes driven by biphasic stimulator (model 305C; Aurora Scientific Inc.). L_o_ determined using single twitches (pulse width=0.2ms) at 1 Hz stimulation frequency with 1 minute rest in between contractions to avoid fatigue. Once L_o_ was determined, the strip acclimatized for 30 minutes in the oxygenated bath. L_o_ was re-assessed and measured with a ruler and the start of the force-frequency protocol was initiated (1, 10, 20, 40, 60, 80, 100, 120, 140 and 200Hz with 1 minute rest in between contractions). Force production was normalized to CSA of the muscle strip (m/l*d) where m is the muscle mass, l is the length, and d is mammalian skeletal muscle density (1.06mg.mm^3^).

### 2.6 Western Blotting

A frozen piece of TA and diaphragm from each animal was homogenized in a plastic microcentrifuge tube with a tapered Teflon pestle in ice-cold buffer containing (in mM) 20 Tris/HCl, 150 NaCl, 1 EDTA, 1 EGTA, 2.5 Na_4_O_7_P_2_, and 1 Na_3_VO_4_ and 1% Triton X-100 with PhosSTOP inhibitor tablet (Roche; 4906845001) and protease inhibitor cocktail (Sigma Aldrich; P8340) (pH7.0) as published previously [16,30]. Protein concentrations were determined using a bicinchoninic acid assay (life Technologies, Thermo Fisher Scientific). 15-20 μg of denatured and reduced protein was subjected to 10%-12% gradient SDS-PAGE followed by transfer to low-fluorescence polyvinylidene difluoride membrane. Membranes were blocked with Odyssey Blocking Buffer (Li-COR) and immunoblotted overnight (4^ο^C) with antibodies specific to each protein. A commercially available monoclonal antibody was used to detect electron transport chain proteins (rodent OXPHOS Cocktail, ab110413; Abcam, Cambridge, UK, 1:250 dilution), including V-ATP5A (55kDa), III-UQCRC2 (48kDa), IV-MTCO1 (40kDa), II-SDHB (30 kDa), and I-NDUFB8 (20 kDa). A commercially available monoclonal antibody was used to detect mitochondrial creatine kinase (mtCK C-1 43 kDa; Santa Cruz 376320, 1:500 dilution).

After overnight incubation in primary antibodies, membranes were washed 3 times for 5 minutes in TBS-Tween and incubated for 1 hour at room temperature with the corresponding infrared fluorescent secondary antibody (LI-COR IRDye 680RD 925-68020, 1:20 000).

### 2.7 Preparation of permeabilized muscle fibers

The assessment of mitochondrial bioenergetics was performed as described previously in our publications [16,23,31–33]. Briefly, the TA and diaphragm from the mouse was removed and placed in ice cold BIOPS. Muscle was separated gently along the longitudinal axis to from bundles that were treated with 40 μg/mL saponin in BIOPS on a rotor for 30 minutes at 4^ο^C. Following permeabilization the permeabilized muscle fiber bundles (PmFBs) for respiration were blotted and weighed in 1.5mL of tared prechilled BIOPS for normalization of respiratory assessments. The remaining PmFBs for mitochondrial H_2_O_2_ (mH_2_O_2_) were not weighed at this step as these data were normalized to fully recovered dry weights taken after the experiments. All PmFBs were then washed in Buffer Z on a rotator for 15 minutes at 4^ο^C to remove the cytoplasm. Buffer Z contained (in mM) 105 K-MES, 30 KCl, 10 KH_2_PO_4_, 5 MgCl_2_·6 H_2_O, 1 EGTA and 5mg/mL BSA (pH 7.1).

### 2.8 Mitochondrial respiration

High-resolution respirometry (O_2_ consumption) were conducted in 2 mL of respiration medium (Buffer Z) using the Oroboros Oxygraph-2k (Oroboros Instruments, Corp., Innsbruck, Austria) with stirring at 750 rpm at 37°C. Buffer Z contained 20 mM Cr to saturate mtCK and promote phosphate shuttling through mtCK or was kept void of Cr to prevent the activation of mtCK [34]. All experiments were conducted in the presence of 5 μM blebbistatin (BLEB) in the assay media to prevent spontaneous contraction of PmFB, which has been shown to occur in response to ADP at 37°C that alters respiration rates [34]. Complex I-supported respiration was stimulated using 5mM pyruvate and 2mM malate (NADH) followed by a titration of ADP concentrations from physiological ranges (25 μM, 100 μM; [35]) to high submaximal (500 μM) and saturating to stimulate maximal coupled respiration (5000 μM in the presence of creatine and 7000 μM in the absence of creatine). 10mM glutamate (further NADH generation) was added at the end of the ADP titration. Cytochrome *c* was then added to test mitochondrial outer membrane integrity. Experiments with low ADP-stimulated respiration (bundles that did not respond to ADP) with high cytochrome *c* responses (>15% increase in respiration) were removed from analysis (13 of 370 bundles). Last, 20mM Succinate (FADH_2_) was added to stimulate complex-II supported respiration. These protocols were designed to understand the regulation of respiration coupled to oxidative phosphorylation of ADP to ATP (Adenosine triphosphate).

### 2.9 mH_2_O_2_

mH_2_O_2_ was determined spectrofluorometrically (QuantaMaster 40, HORIBA Scientific) in PmfB placed in a quartz cuvette with continuous stirring at 37°C in 1 mL of Buffer Z supplemented with 10 μM Amplex Ultra Red, 0.5 U/ml horseradish peroxidase, 1mM EGTA, 40 U/ml Cu/Zn-SOD1, 5 μM BLEB and 20mM Cr to saturate mtCK. No comparisons were made to PmFB in the absence of creatine due to tissue limitations. State II mH_2_O_2_ (maximal emission in the absence of ADP) was induced using the Complex I-supporting substrates (NADH) pyruvate (10mM) and malate (2mM) mH_2_O_2_ to generate forward electron transfer (FET)-supported electron slip at Complex I [36] as described previously [23]. These PmFBs were incubated with 35 μM CDNB during the 30-minute permeabilization to deplete glutathione and allow for detectable rates of mH_2_O_2_. Following the induction of State II mH_2_O_2_, a titration of ADP was employed to progressively attenuate mH_2_O_2_ as it occurs when membrane potential declines during oxidative phosphorylation [37]. A separate PmFB was used to stimulate electron slip at Complex I through reverse electron transfer (RET) from complex II using succinate (FADH_2_) [36] followed by ADP titrations as used in the previous protocol. After the experiments, the fibres were lyophilized in a freeze-dryer (Labconco, Kansas City, MO, USA) for > 4h and weighed on a microbalance (Sartorius Cubis Microbalance, Gottingen Germany). The rate of mH_2_O_2_ emission was calculated from the slope (F/min) using a standard curve established with the same reaction conditions and normalized to fibre bundle dry weight.

### 2.10 Serum GDF15

Blood was collected through cardiac puncture and allowed to clot at room temperature for 30 minutes. Blood was then spun at 1000g for 10 minutes and serum was collected. GDF15 (Growth differentiation factor 15) levels were analyzed in serum using the mouse GDF-15 DuoSet ELISA kit according to the manufacturer’s instructions (R&D Systems DY6385).

### 2.11 RNA isolation and Rt-PCR

To perform reverse transcription-polymerase chain reaction (Rt-PCR), tissue was used from a separate cohort of mice. These mice were ID8-inoculated as done previously and housed at the University of Guelph. These mice had cancer for similar times (∼45, ∼75, ∼90 days), and mice at the 45-day time point were 15-16 weeks old at the time of euthanasia. RNA isolation was performed twice for two separate analyses.

In the first analysis, RNA isolation was performed at the University of Guelph using a TRIzol (Invitrogen) and RNeasy (Qiagen) hybrid protocol. Briefly, snap frozen TA tissue was homogenized in 1mL of TRIzol reagent according to the manufacturer instructions. The RNA mixture was transferred to a RNeasy spin column (Qiagen) and processed according to the RNeasy kit instructions. RNA was quantified spectrophotometrically at 260nm using a NanoDrop (ND1000, ThermoFisher Scientific INC.) and used for RNA sequencing.

In the second analysis, RNA isolation was performed at York University on a separate cohort of tissue as previously described [29]. TA and diaphragm samples were lysed using TRIzol reagent (Invitrogen) and RNA was separated to an aqueous phase using chloroform. The aqueous layer containing RNA was then mixed with isopropanol and loaded to Aurum Total RNA Mini Kit columns (Bio-Rad, Mississauga, ON, Canada). Total RNA was then extracted according to the manufacturer’s instructions. RNA was quantified spectrophotometrically using the NanoDrop attachment for the Varioskan LUX Multimode Microplate reader (Thermo Scientific). Reverse transcription of RNA into cDNA was performed by M-MLV reverse transcriptase and oligo(dT) primers (Qiagen, Toronto, ON, Canada). cDNA was then amplified using aCFX384 Touch Real-Time PCR Detection Systems (Bio-Rad) with a SYBR Green master mix and specific primers **(STable 1)**. Gene expression was normalized to β-actin (Actb) and relative differences were determined using the ΔΔCt method. Values are presented as fold changes relative to the control group.

### 2.12 RNA Sequencing

RNA libraries were prepared using the NEBNext Ultra II Directional RNA Library Prep Kit for Illumina (NEB, E7760) according to manufacturer’s polyA mRNA workflow at the Advanced Analysis Center at the University of Guelph (Guelph, Ontario, Canada). Libraries were normalized, denatured, diluted, and sequenced on an Illumina 2×100 bp NovaSeq S4 flowcell usingv1.5 chemistry according to manufacturer’s instructions.

After demultiplexing, fastq files were uploaded into Partek and pre- and post-alignment quality control (QC) was performed in Partek (average phred quality score >35). Paired-end reads (100bp or 150bp) were aligned using STAR 2.7.8a [38] with mm39 – RefSeq Transcripts 98 (05/05/2021) in Partek. A minimum read cutoff of 20 was applied; all other settings were default. Data were normalized using counts per million (CPM), and Limma-voom [39] was used for differential gene expression analysis: sham vs 45d, sham vs 75d, and sham vs 90d. Raw p-values were adjusted for false discovery rate (FDR) using the FDR step-up procedure. Gene Ontology and Reactome pathway enrichment analysis was completed using Enrichr and ConsensusPathDB with a background list, using DEGs with *p* < 0.05. Significance threshold for volcano plots were set at P <0.01.

### 2.13 Statistics

Results are expressed as mean ± SD. The level of significance was established at *p* <0.05 for all statistics. The D’Agostino-Pearson omnibus normality test was first performed to determine whether data resembled a Gaussian distribution, and all data were subject to the ROUT test (Q=0.5%) to identify and exclude outliers which was a rare occurrence. When data fit normal distributions, standard one way and two-way ANOVAs were performed. When data did not fit a Gaussian distribution for analysis with one independent variable, the Kruskal-Wallis test was used. Moreover, when data did not fit a Gaussian distribution for analysis with two independent variables, data was first log transformed then analyzed using a standard two-way ANOVA (See **SFigure 7** for log transformed analysis of **Figure 6A, 6B, 6E, 7D, 7F, 7I and 7K**) but data was still presented in the Results as non-transformed data. Respective statistical tests are provided in figure legends. When significance was observed with an ANOVA, post-hoc analyses were performed with a two-stage set-up method of Benjamini, Krieger and Yekutieli for controlling false discovery rate (FDR) for multiple-group comparisons. With this method, all reported *p* values are FDR adjusted (Traditionally termed “q”). All statistical analyses were performed in GraphPad Prism 10 (La Jolla, CA, USA).

## 3. Results

### 3.1 Orthotopic epithelial ovarian cancer induces metastasis, ascites, impaired functional capacity and body weight loss at advanced stages

Immunocompetent C57BL6 mice received orthotopic injections of murine ID8 epithelial ovarian cancer cells **(Figure 1A)**. Tumours were allowed to develop for ∼45 days, ∼75 days and ∼90days post tumour injection to evaluate muscle responses across tumour development. This study was completed in two cohorts of mice to obtain sufficient tissue to complete all experiments. Weekly body weights (BW) in both cohorts were tracked post tumour injection **(Figure S1A and S1B).** Body weight, tumour weight, ascitic volume, muscle mass and spleen mass data were then merged as these data were collected in both cohorts.

Primary ovarian tumour mass grew progressively, reaching a maximum of ∼400mg by ∼90 days **(Figure 1B)**. Metastatic tumour spread to the diaphragm was noted at the 90 day timepoint **(Figure 1C)** observed with abundant mononuclear cell infiltration **(Figure 1D and 1E).** Another common secondary complication in ovarian cancer patients is the development of ascites fluid within the intraperitoneal space. Ascites developed as early as 77 days post ID8-inoculation (data not shown); and there was a significant increase in the amount of ascites paracentesis taps performed and volume of ascites collected by 90 days post-inoculation **(Figure 1F-H)**. At this time, decreases in voluntary wheel running and grip strength were noted **(Figure 1I and 1J).** Change in weekly body weight (BW), tibia length and peak body mass demonstrate how all groups grew similarly post cancer injection, as there were no significant differences between groups (**Figure 1K-1M)**. However, final primary tumour-free BW and % change of peak BW to final BW were significantly decreased in the 90-day group indicative of cachexia **(Figure 1N & 1O)**.

### 3.2 Muscle loss and fat loss occur at advanced stages of ovarian cancer with no increases in GDF15, inflammatory markers or atrogenes

In addition to body weight loss, muscle mass and fat mass loss are hallmarks of cancer cachexia. Muscle mass was lower at the 90-day time point in the extensor digitorum longus (EDL; −11%), plantaris (PLT; −18%), tibialis anterior (TA; −19%), gastrocnemius (GA; −13%), and quadriceps (Quad; −13%) muscles compared to control (CTRL) whereas Soleus (SOL) mass did not change **(Figure 2A).** Adipose tissue from the inguinal fat pad was lower in the 45 Day and 90 Day groups post ovarian cancer inoculation **(Figure 2B).** Serum GDF15 – a recently identified cachexia regulator [40] – did not change **(Figure 2 C).** Spleen mass was greater in the 75 and 90 Day groups suggesting an increased inflammatory response **(Figure 2D)**. We then measured cytokines and atrophy markers known to be elevated in certain clinical and preclinical models of cancer cachexia [40,41]. In the TA, Interleukin 6 (IL-6) mRNA was not different between groups while tumour necrosis factor alpha (TNF-α) was lower at 45 days compared to control **(Figure 2E).** In addition, atrogin and muscle RING-finger protein-1 (MURF-1) mRNA followed similar patterns whereby mRNA levels were lower at early time points with no differences compared to control in the 90 Day group **(Figure 2F)**. In the diaphragm, IL-6 mRNA was higher in the 45 and 90 Day groups, while TNF-α was lower at the 45 and 75 Day groups **(Figure 2G).** Atrogin mRNA was lower than control in the 90 Day group while MURF-1 was not different between groups **(Figure 2H)**.

### 3.3 Muscle atrophy in the absence of muscle regeneration occurs earlier in the diaphragm compared to TA throughout ovarian cancer development

Muscle atrophy is also a hallmark of cancer cachexia. TA and diaphragm muscles were sectioned and tagged for MHC isoforms I, IIA and IIB; black fibers were assumed IIx. In the TA, fiber CSA was not different in the 45 Day group in any isoform or when isoforms were pooled **(Figure 3A-C**). In the 75 Day group, TA exhibited lower CSA in pooled fibers (−11%) with specific reductions in MHCIIx isoforms **(Figure 3A-C).** In the 90 Day group, TA exhibited an exacerbated reduction in pooled fiber CSA (−23%) with specific reductions in all isoforms compared to control **(Figure 3A-C).** This is further exemplified when pooled fibers are presented in a histogram as a higher frequency of smaller fibers exist in the TA in the 90 Day group compared to control **(Figure 3D)**. We also demonstrate muscle atrophy was occurring in the absence of apparent regeneration by measuring eMHC to evaluate if new fibers were developing. In the TA no eMHC fibers were identified in any groups **(Figure 3E and 3F)**.

**Figure 3.**
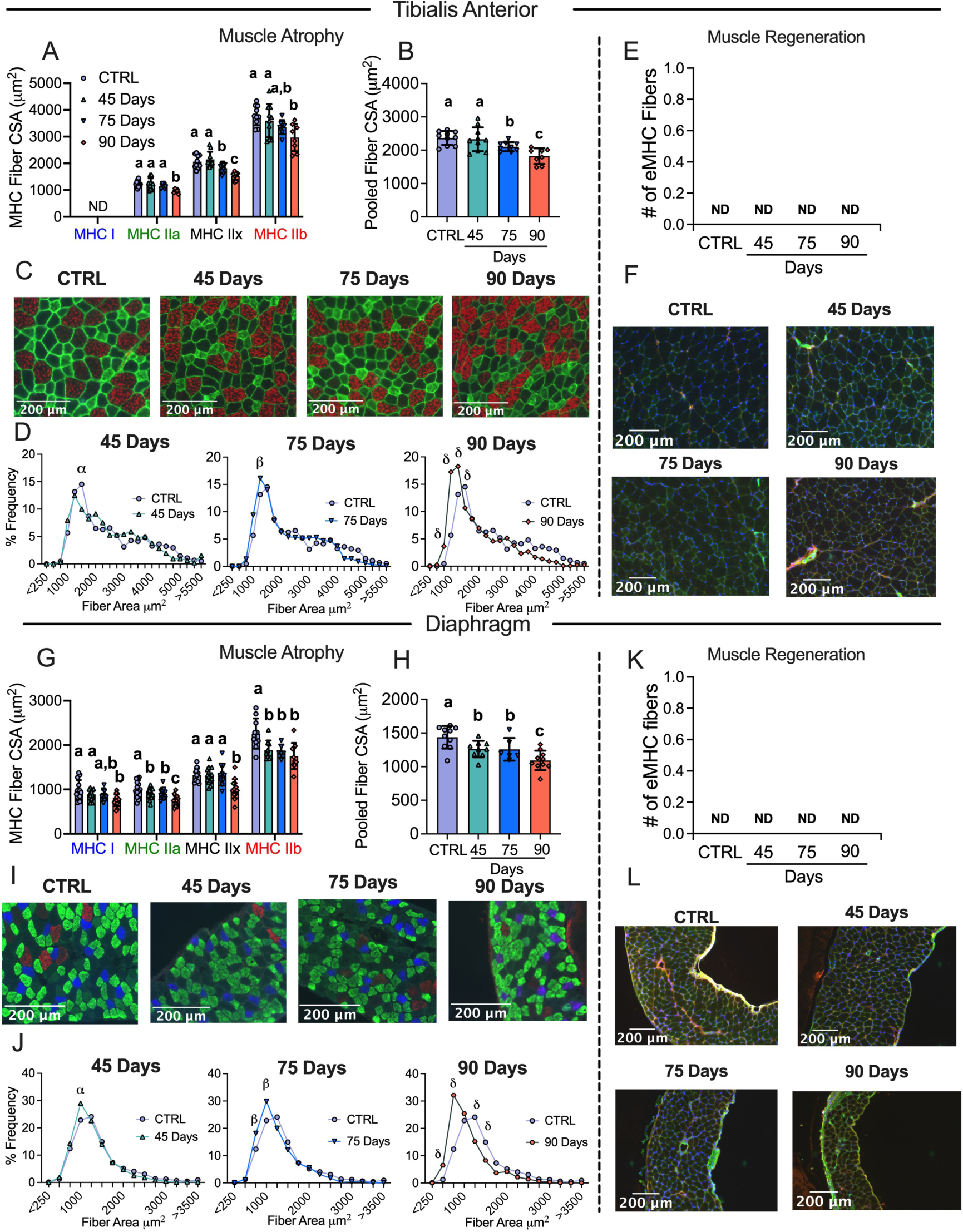
Evaluation of tibialis anterior and diaphragm fiber type atrophy and fiber regeneration in epithelial ovarian cancer injected mice. Analysis of fiber histology on myosin heavy chain (MHC) isoforms and eMHC was performed in control and EOC mice. Cross-sectional area (CSA) of MHC isoforms were evaluated in the tibialis anterior (A-C, n = 7-10). All fiber types were also pooled, binned and averaged based off fiber area and plotted by frequency distribution at each time point compared to control (D, n =7-10). Embryonic MHC (eMHC) was tagged in separate sections to evaluate the presence of new fibers (E & F, n = 7-10). This was repeated within the diaphragm (G-L, n 10 = 14). Results represent mean ± SD. Lettering denotes statistical significance when different from each other (*p* < 0.05). α *p* < 0.05 Control versus 45 Days; β *p* < 0.05 Control versus 75 Days; δ *p* < 0.05 Control versus 90 Days. A one-way ANOVA was used for figures A, B, G and H. Data that was not normally distributed was analyzed with a Kruskal-Wallis test. A two-way ANOVA was used for figures D and J (interactions shown only). All ANOVAs were followed by a two-stage step-up method of Benjamini, Krieger and Yukutieli multiple comparisons test. C57BL/6J female mice ∼75 days post PBS injection as controls (CTRL); C57BL/6J female mice ∼45 days post ovarian cancer injection (45 Days); C57BL/6J female mice ∼75 days post ovarian cancer injection (75 Days); C57BL/6J female mice ∼90 days post ovarian cancer injection (90 Days).

In the diaphragm, muscle atrophy was evident earlier than in the TA. In the 45 Day group, diaphragm fibers exhibited lower CSA stained with MHCIIa and MHCIIb isoforms as well as lower CSA in pooled fibers (−12%) **(Figure 3G-I)**. In the 75 Day group, fiber CSA remained lower in pooled fibers (−13%) specifically in MHCIIa and MHCIIb isoforms compared to control and were similar to the 45 Day group **(Figure 3G-I).** In the 90 Day group, diaphragm CSA exhibited extensive reductions when isoforms were assessed separately or pooled (−24%) **(Figure 3G-I).** This is further demonstrated when pooled fibers are displayed in a histogram as a higher frequency of smaller fibers were present at 90 days **(Figure 3J).** This atrophy in the diaphragm also occurred in the absence of apparent regeneration as no eMHC fibers were identified **(Figure 3K and 3L).**

### 3.4 Early muscle weakness is further reduced in the diaphragm throughout ovarian cancer progression but gradually recovers in the TA

Muscle weakness is another hallmark of cancer cachexia. In the 45 Day group, TA specific force production was lower compared to control **(Figure 4A and 4B)**. In contrast, force production increased progressively by 75 days and 90 days, while still remaining lower compared to control (main effect) **(Figure 4A and 4B)**. The rate of contraction (Df/dt) at 1Hz stimulation frequency was lower at 45 Day group compared to control but not different at 100Hz **(Figure 4C)**. In addition, the half relaxation time (HRT) was significantly longer in the 90 Day group at both stimulation frequencies **(Figure 4D)**. This suggests that at 45 days the TA exhibits a slower rate of contraction at lower stimulation frequencies, and at 90 days the TA exhibits slower relaxation. We also measured mRNA content of ryanodine receptors (RyR1) and sarcoendoplasmic reticulum calcium ATPase (SERCA) as these proteins are integral for the regulation of calcium release and reuptake that regulates contraction. RyR1 mRNA content was not different across the time points, however, SERCA1 mRNA content was higher in the 90 Day group compared to control **(Figure 4E)**.

**Figure 4.**
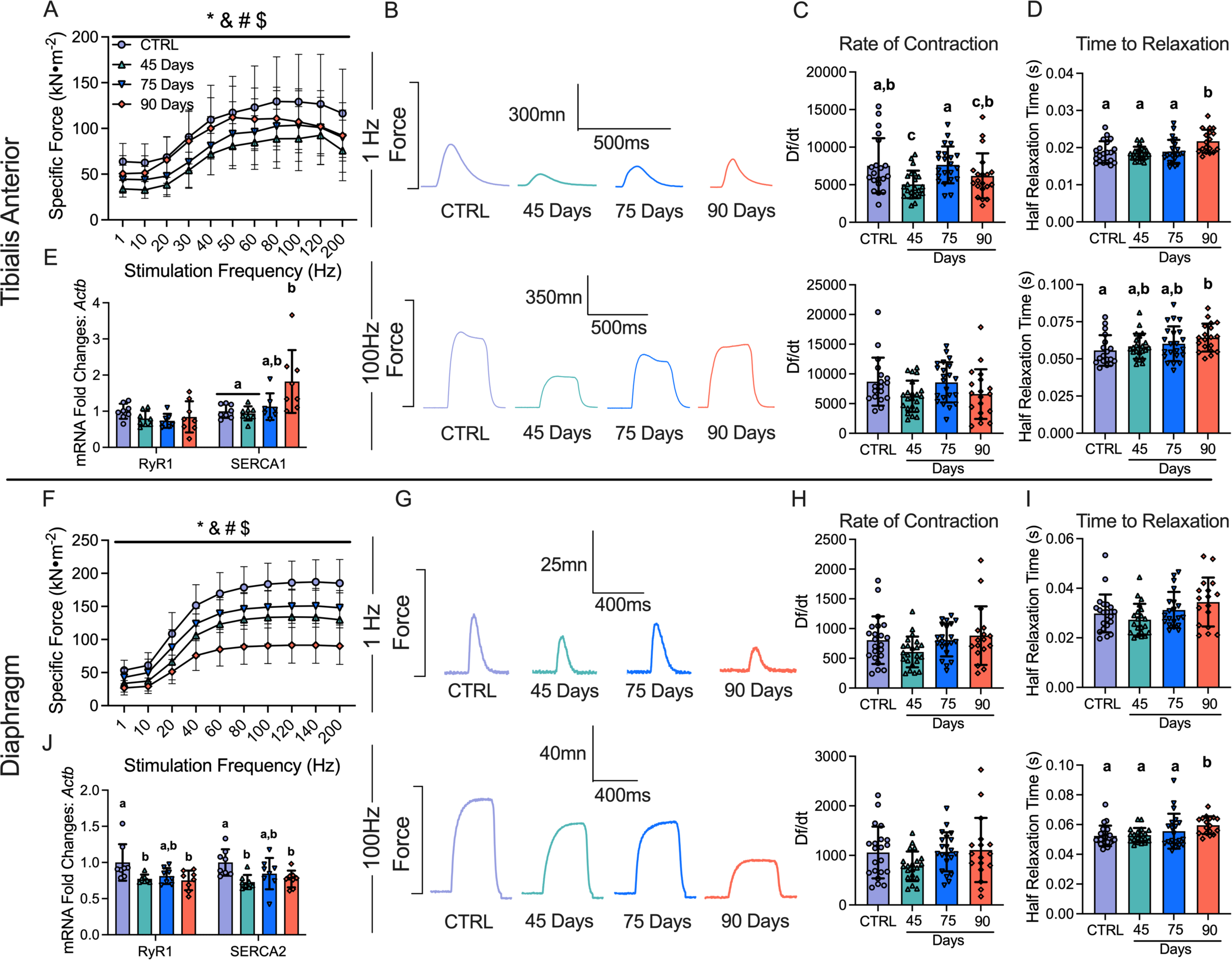
The effects of epithelial ovarian cancer (EOC) on tibialis anterior and diaphragm force production, contractile properties and calcium handling gene expression. In situ tibialis anterior force production was assessed using the force-frequency relationship (A, n = 9-10; B, Representative twitches at 1 Hz and 100Hz). Rate of twitch contractions along with the half relaxation time were also assessed at 1Hz and 100Hz (C & D, n = 18-22). mRNA expression of ryanodine receptors (RyR1) and sarcoplasmic/endoplasmic reticulum ATPase (SERCA; SERCA1 used for tibialis anterior (fast twitch) and SERCA2 for diaphragm (slow twitch) was also measured (E, n = 8). This was repeated for the diaphragm (F-J, n = 8-22) Results represent mean ± SD. * *p* < 0.05 Control versus all time points; & *p* < 0.05 45 Days versus all time points; # *p* < 0.05 75 Days versus all time points; $ *p* < 0.05 90 Days versus all time points. Lettering denotes statistical significance at an alpha set at *p* < 0.05. A two-way ANOVA was used for figures A and J (main effects shown only) and all other data was analyzed using a one-way ANOVA or Kruskal-Wallis test when data did not fit normality. All ANOVAs were followed by a two-stage step-up method of Benjamini, Krieger and Yukutieli multiple comparisons test. C57BL/6J female mice ∼75 days post PBS injection as controls (CTRL); C57BL/6J female mice ∼45 days post ovarian cancer injection (45 Days); C57BL/6J female mice ∼75 days post ovarian cancer injection (75 Days); C57BL/6J female mice ∼90 days post ovarian cancer injection (90 Days).

Diaphragm force production was also significantly lower compared to control in the 45 Day group as a main effect (interaction at 40Hz onward; not shown), with no changes in contractile properties but decreases in RyR1 and SERCA2 mRNA contents **(Figure 4F-4J).** Interestingly, in the 75 Day group, diaphragm specific force transiently increased compared to the 45 Day group while still remaining lower than control as a main effect (interaction at 40Hz onward; not shown), with no changes in the rate of contraction, time to relaxation or mRNA content of RyR1 or SERCA2 **(Figure 4F-4J)**. In the 90 Day group, force production was further lowered compared to all time points as a main effect (interaction at 40Hz onward; not shown) with longer half relaxation time at 100Hz stimulation frequency **(Figure 4F-4I)**. This decrease in force production and increase in half relaxation time was coupled to decreases in RyR1 and SERCA2 mRNA content **(Figure 4J)**. These data demonstrate that each muscle demonstrates unique contractile adaptations to ovarian cancer over time.

### 3.5 Mitochondrial genes are downregulated in the TA during early and advanced stages of epithelial ovarian cancer

Given there were no changes in markers of atrophy-related mechanisms that have been found in other cachexia models (GDF15, TNF-α, Atrogin and MURF-1) [40,41], we examined the potential role for mitochondrial stress responses that have been identified in other models [16,42]. In the TA, RNA-seq revealed 691, 795 and 3402 differentially expressed genes (DEGs) in the 45, 75, and 90 Day groups compared to control, respectively. Some of these DEGs are shared among time points **(Figure 5A)**. We then used a volcano plot to demonstrate DEGs that were upregulated or downregulated compared to control **(Figure 5B – 5D).** Gene ontology (GO) enrichment analyses was then performed on DEGs to investigate up and down regulated biological processes altered across time points compared to control **(Figure 5E-5G)**. Several pathways were significantly enriched, with the majority of downregulated pathways being mitochondrial-related. The two most significantly different upregulated and downregulated pathways from the enrichment analysis were represented in a chord plot to exemplify the specific genes changing across ovarian cancer progression **(Figure 5H-5J)**. These results largely demonstrate how more mitochondrial genes are down regulated as ovarian cancer progresses with increases in certain muscle contraction related genes in the 90 Day group **(Figure 5H-5J)**.

**Figure 5.**
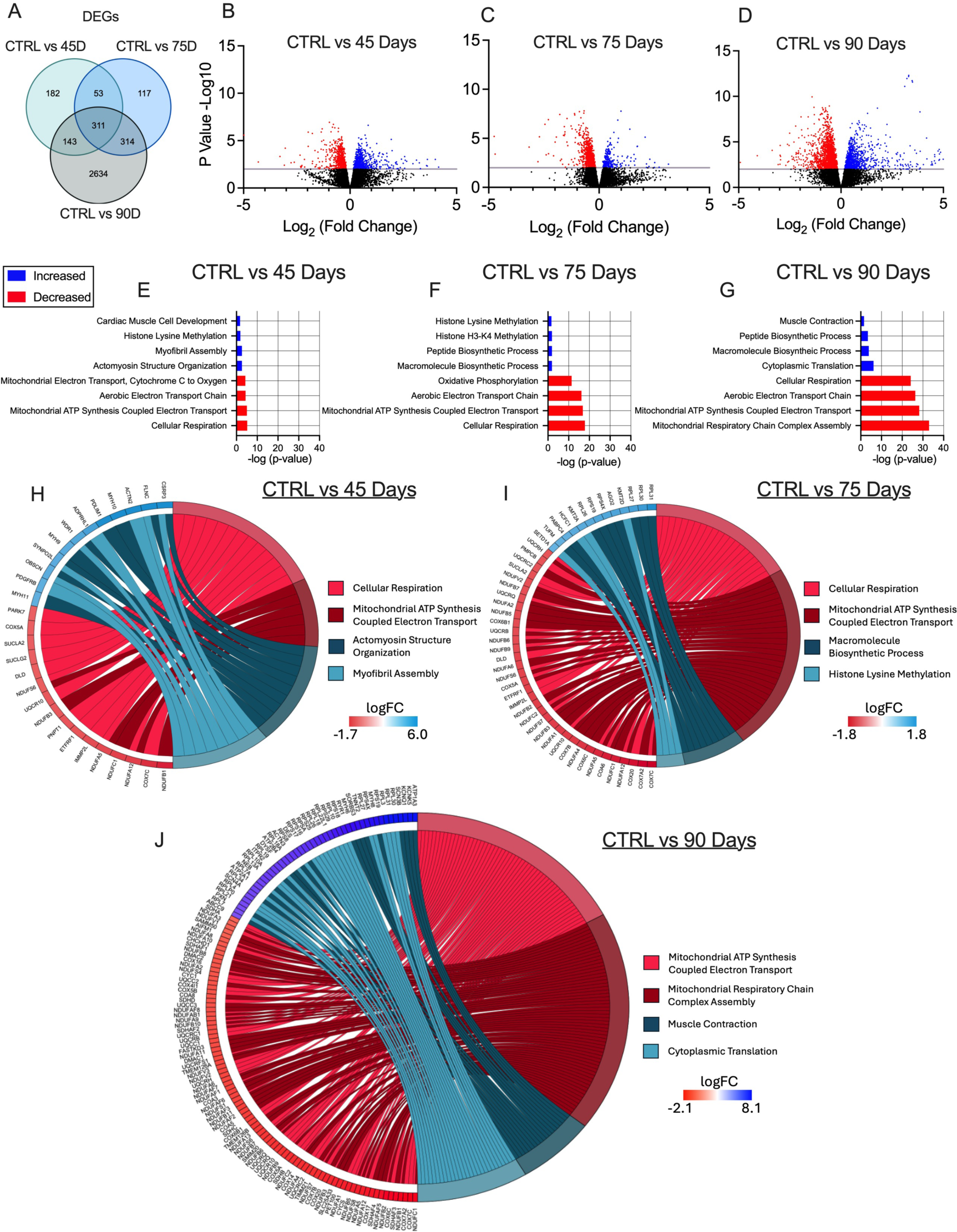
RNA sequencing analysis of tibialis anterior muscle in epithelial ovarian cancer (EOC) injected mice. Number of differentially expressed genes (DEGs) in each comparison were exemplified in a Venn diagram (A). Volcano plot showing −log10 p-value and log2 fold changes of DEGs for each comparison were also completed (B-D). Top 3 upregulated and top 3 downregulated biological processes enriched in DEGs at each time point were also analyzed and graphed (E-G) Top two upregulated and down regulated biological processes were also used to generate a chord plot with the corresponding DEGs and respective log fold changes at each time point (H-J). n = 6. C57BL/6J female mice ∼75 days post PBS injection as controls (CTRL); C57BL/6J female mice ∼45 days post ovarian cancer injection (45 Days); C57BL/6J female mice ∼75 days post ovarian cancer injection (75 Days); C57BL/6J female mice ∼90 days post ovarian cancer injection (90 Days).

### 3.6 Decreases in carbohydrate supported mitochondrial respiration occur in early stages of ovarian cancer progression but is restored by late-stage disease

Using permeabilized muscle fibres, mitochondrial respiration was assessed by stimulating complex I with NADH generated by pyruvate (5mM; carbohydrate substrate) and malate (2mM) across a range of ADP concentrations to challenge mitochondria over a spectrum of metabolic demands. The ADP titrations were repeated without (−Creatine; **Figure 6A & 6D**) and with creatine (+Creatine; **Figure 6B & 6E**) in the assay media to model the two main theoretical mechanisms of high energy phosphate shuttling from the mitochondria to the cytosol [37,43–45]. Briefly, in the absence of creatine, ATP is exported across the double membranes while in the presence of creatine, matrix-derived ATP crosses the inner membrane and is used by mitochondrial creatine kinase in the intermembrane space to phosphorylate creatine. The phosphocreatine product is then exported across the outer membrane which is then used by cytosolic creatine kinases to re-phosphorylate ADP local to ATP consuming proteins. Previous studies have shown that ADP/ATP flux is much slower than creatine/phosphocreatine due to the diffusion limitations of ATP/ADP vs phosphocreatine/creatine [46]. Prior modeling experiments have posited that up to 80% of the phosphate exchange between mitochondria and cytoplasm (in muscle) is likely comprised of the creatine-dependent system. 25μM, 100μM and 500μM ADP were selected to reflect creatine sensitivity as this is within the predicted range that is sensitive to the effects of mitochondrial creatine kinase (mtCK) [23,46–48]. Maximal ADP-stimulated respiration was unchanged in both muscles, in each condition, at all-time points **(SFigure 4A-D)** indicating that cancer alters the regulation of mitochondrial pyruvate oxidation stimulated by submaximal ADP concentrations.

**Figure 6.**
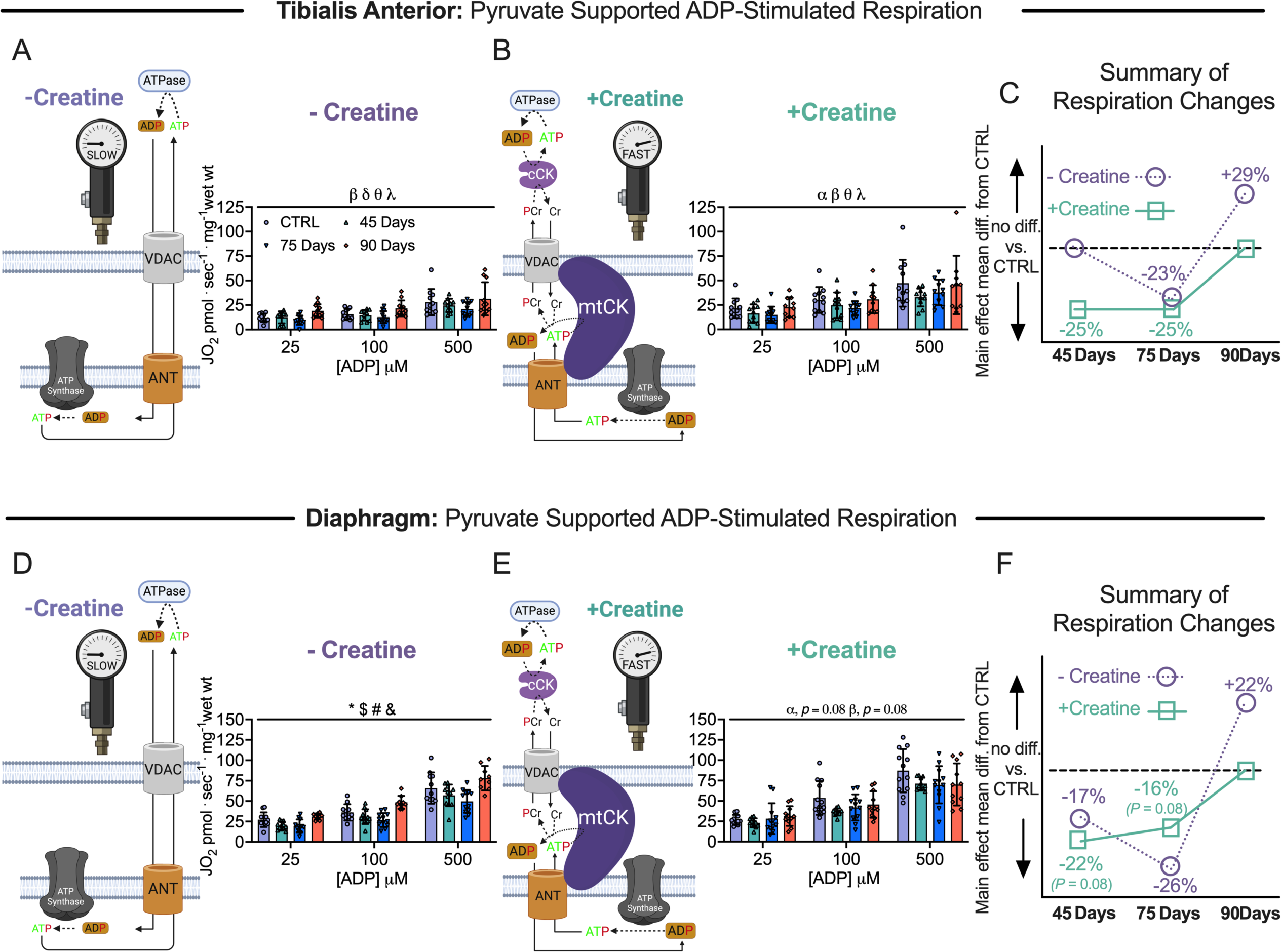
Pyruvate & malate supported mitochondrial respiration in tibialis anterior and diaphragm muscle of epithelial ovarian cancer (EOC) injected mice. ADP-stimulated (State III) respiration supported by pyruvate (5mM) and malate (2mM) generating NADH was assessed in the absence (−creatine) and presence (+creatine) of creatine within tibialis anterior and diaphragm PmFBs of EOC injected mice. Mitochondrial respiration in the absence of creatine was assessed at submaximal concentrations (25μM, 100μM and 500μM) in tibialis anterior of EOC injected (A). Mitochondrial respiration in the presence of creatine was also assessed at submaximal concentrations (25μM, 100μM and 500μM) in tibialis anterior of EOC injected (B). A schematic representative summary of changes in -Creatine/+Creatine pathways is depicted (C). This was repeated for the diaphragm (D-F) Results represent mean ± SD. n = 9-12. α *p* < 0.05 Control versus 45 Day; β *p* < 0.05 Control versus 75 Day; δ *p* < 0.05 Control versus 90 Day; θ *p* < 0.05 45 Day versus 90 Day; λ *p* < 0.05 75 Day vs 90 Day; * *p* < 0.05 Control versus all time points; & *p* < 0.05 45 Days versus all time points; # *p* < 0.05 75 Days versus all time points; $ *p* < 0.05 90 Days versus all time points. All ANOVAs were followed by a two-stage step-up method of Benjamini, Krieger and Yukutieli multiple comparisons test. Voltage dependent anion channel (VDAC); adenine nucleotide translocator (ANT); mitochondrial creatine kinase (mtCK); adenosine diphosphate (ADP); adenosine triphosphate (ATP); phosphocreatine (PCr); creatine (Cr); creatine-independent phosphate shuttling (−Creatine); creatine-dependent phosphate shuttling (+Creatine). All data was analyzed suing a two-way ANOVA (main effects shown only). C57BL/6J female mice ∼75 days post PBS injection as controls (CTRL); C57BL/6J female mice ∼45 days post ovarian cancer injection (45 Days); C57BL/6J female mice ∼75 days post ovarian cancer injection (75 Days); C57BL/6J female mice ∼90 days post ovarian cancer injection (90 Days).

Within the 45 and 75 Day groups, TA and diaphragm exhibit a decrease in mitochondrial respiration in the -Creatine condition with an apparent supercompensation by 90 days **(Figure 6A & 6D)**. In the +Creatine condition, early decreases in mitochondrial respiration in both muscles returned to normal by the 90-day time point **(Figure 6B & 6E)**. A summary of all respiration changes across time and between conditions as a main effect versus control is provided for both muscles **(Figure 6C & 6F).** All changes in mitochondrial respiratory control were not influenced by changes in mitochondrial electron transport chain (ETC) content as there were no changes in ETC subunit content estimated via western blot **(Figure S3A & S3B).**

Another approach for evaluating changes in the relative control exerted by mitochondrial creatine kinase on ADP-stimulated respiration, or ‘mitochondrial creatine sensitivity’, is to calculate the ratio of respiration at submaximal ADP concentrations in both +Creatine/-Creatine conditions [16,23,28,46,49]. In the TA, creatine sensitivities were unchanged across time points indicating alterations in respiration were similar between both high energy phosphate shuttling systems **(SFigure 4E)**. While both -Creatine and +Creatine respiration changed across time points, the relative changes were disproportionate within the diaphragm, particularly in the 90 Day group compared to control. More specifically, the -Creatine system exhibited a greater increase in respiration compared wo the +Creatine system, thus, mitochondrial sensitivity to creatine was decreased compared to control in the 90 Day group, but this does not necessarily reflect compromised energy transfer from mitochondria to cytoplasm given both systems nonetheless improved. **(SFigure 4F)**, indicating the creatine-dependent phosphate shuttling system is selectively impaired compared to the creatine-independent system. This effect was not explained by mtCK protein contents given they were unchanged at all time points **(SFigure 4G & 4H).**

We also measured fat oxidation in each muscle in order to determine if these changes in respiration were unique to pyruvate stimulation of respiration through Complex I (NADH). In the TA, there were no changes in palmitoyl CoA-supported respiration (**SFigure 5B & 5D**) suggesting that the decreases in pyruvate oxidation were not due to a dysfunction that impacted all substrates. Likewise, there were no changes in Complex II-supported respiration (succinate; FADH_2_) and glutamate-supported respiration (further NADH generation; Complex I) in both +Creatine/- Creatine conditions **(SFigure 6A-D)**. This suggests that the decrease in pyruvate oxidation were also not due to Complex I impairments per se nor ETC components downstream of both Complex I and II (see discussion). Interestingly, there was an increase in state II pyruvate/malate respiration in the 90 Day group in the absence of creatine suggesting increased proton leak, but no changes were observed in the presence of creatine **(SFigure 6E & 6F)** nor in response to palmitoyl CoA (**SFigure 5A**).

The diaphragm also exhibited no changes in palmitoyl CoA (**SFigure 5C & 5D**) respiration. However, Complex II-supported respiration (succinate; FADH_2_) in the -Creatine condition was higher in the 90 day group along with glutamate-supported respiration (further NADH generation; Complex I) in both creatine conditions (**SFigure 6G-J**). There were no changes in state II pyruvate/malate-supported respiration **(SFigure 6K & 6L).**

### 3.7 Increased mitochondrial H_2_O_2_ emissions occur in complex I forward and reverse electron transfer in this EOC model

We stimulated complex I-supported mH_2_O_2_ with forward electron transfer (pyruvate and malate (2mM) to generate NADH) **(Figure 7A)** and reverse electron transfer (succinate to generate FADH_2_) **(Figure 7B)** [50–53]. These substrate-specific maximal mH_2_O_2_ kinetics were followed by titration of ADP to determine the ability of ADP to attenuate mH_2_O_2_ during oxidative phosphorylation (OXPHOS). In the TA, pyruvate/malate and succinate supported maximal mH_2_O_2_ was not different at any time point compared to control **(Figure 7C & 7E)**. However, pyruvate/malate supported mH_2_O_2_ during OXPHOS were increased in the 75 Day group which returned to control levels by the 90 Day group, while succinate supported mH_2_O_2_ was increased at the 75 and 90 Day groups **(Figure 7D & 7F).** There were no changes in diaphragm pyruvate/malate and succinate supported maximal mH_2_O_2_ at any time point **(Figure 7H & 7J).** The diaphragm exhibited no changes in pyruvate & malate-supported H_2_O_2_ during OXPHOS but exhibited higher succinate-supported mH_2_O_2_ during OXPHOS in the 75 Day group that returns to baseline by the 90 Day group **(Figure 7I & 7K)**. A summary of mH_2_O_2_ changes across time and between conditions as a main effect versus control is provided for the TA **(Figure 7G)** and diaphragm **(Figure 7L)**.

**Figure 7.**
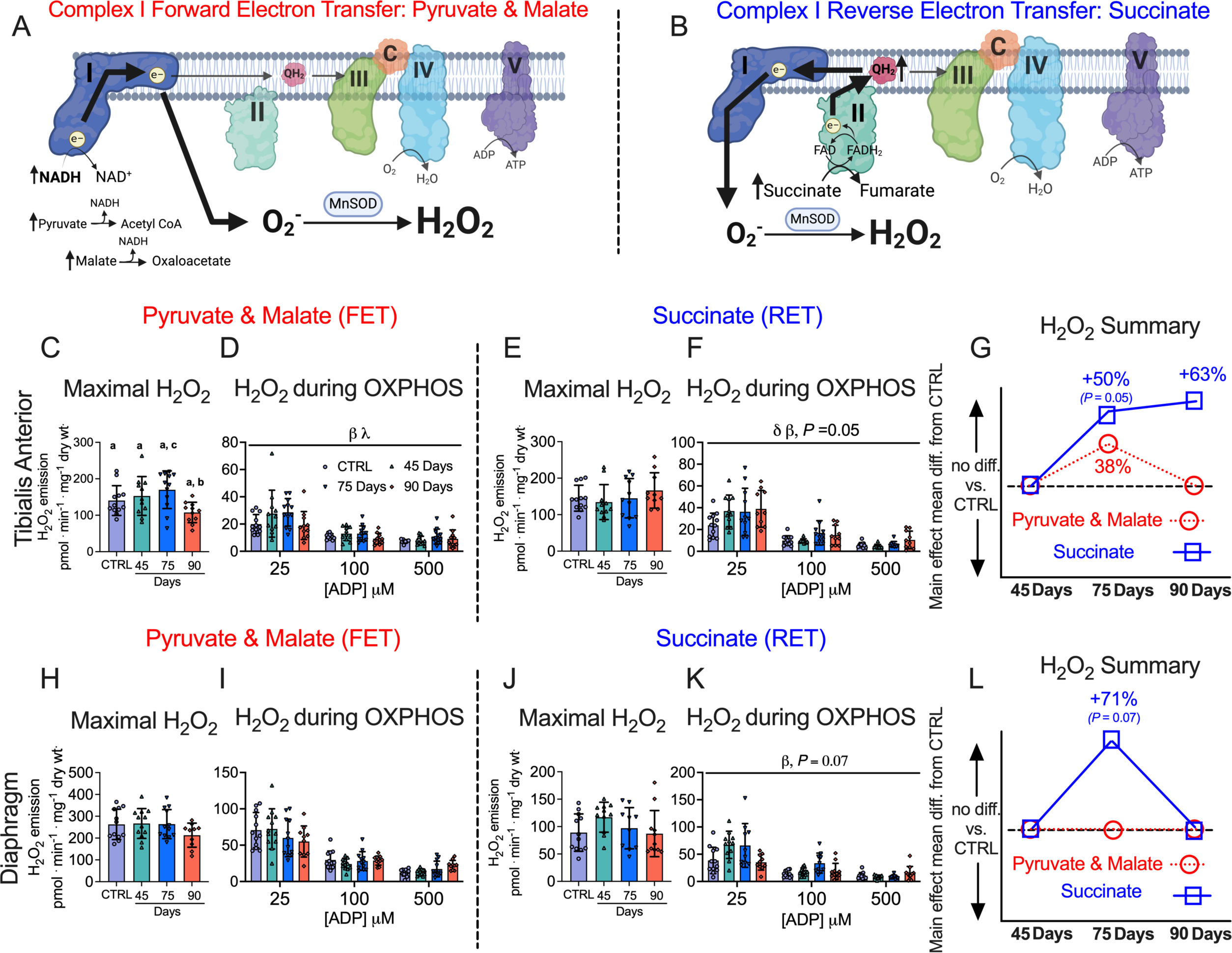
Complex I forward and reverse electron transfer emissions in tibialis anterior and diaphragm muscle of epithelial ovarian cancer (EOC) injected mice. Complex I forward electron transfer (FET) and complex I reverse electron transfer (RET) is schematically depicted (A & B). In FET mitochondrial H_2_O_2_ emission was supported by pyruvate (10mM) and malate (2mM) to generate maximal rates and with ADP to assess H_2_O_2_ emission during OXPHOS. This experiment was repeated to assess RET H_2_O_2_ emission by using succinate (10mM) as opposed to pyruvate and malate. FET and RET H_2_O_2_ emissions were assessed in the TA of EOC injected mice and a summary of changes compared to control is depicted (C-G). This was repeated in the diaphragm (H-L). Results represent mean ± SD. Lettering denotes statistical significance when different from each other (*p* < 0.05). β *p* < 0.05 Control versus 75 Day; λ *p* < 0.05 75 Day vs 90 Day; δ *p* < 0.05 Control versus 90 Day. A one-way ANOVA or Kruskal-Wallis test was used when data did not fit normality in figure C, E, H and J. A two-way ANOVA was used in figured D, F, I and K. All ANOVAs were followed by a two-stage step-up method of Benjamini, Krieger and Yukutieli multiple comparisons test. Oxidative phosphorylation (OXPHOS); manganese superoxide dismutase (MnSOD); electron (e-); superoxide (O ^-^). C57BL/6J female mice ∼75 days post PBS injection as controls (CTRL); C57BL/6J female mice ∼45 days post ovarian cancer injection (45 Days); C57BL/6J female mice ∼75 days post ovarian cancer injection (75 Days); C57BL/6J female mice ∼90 days post ovarian cancer injection (90 Days).

A comprehensive summary of all changes between the TA and diaphragm captured within this study design is provided **(Figure 8).**

**Figure 8.**
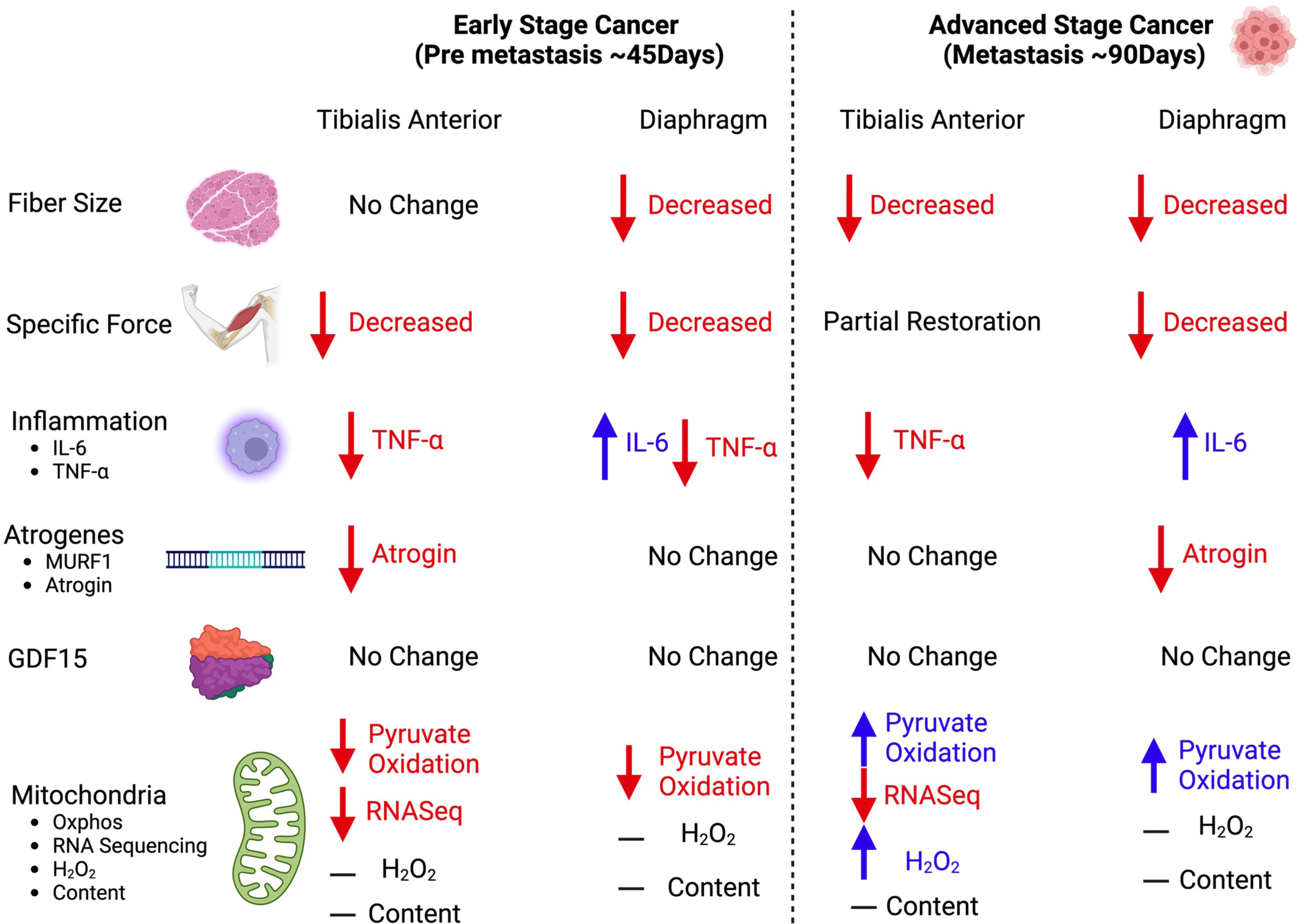
Summary of changes in a metastatic epithelial ovarian cancer cachexia model. When mice were injected with epithelial ovarian cancer, at a pre-metastasis time point (45 days post-injection) early muscle weakness was associated with decreases in pyruvate oxidation in both the tibialis anterior and diaphragm muscles. At this time, the tibialis anterior muscle did not exhibit muscle atrophy while the diaphragm did. With the exception of IL-6 in the diaphragm, there were no increases in TNF-*α* and atrophy markers of cancer cachexia at this time. During robust metastasis (90 days post-injection) both muscles exhibited muscle atrophy and muscle weakness, however the tibialis anterior recovered specific force production. Moreover, both muscles exhibited compensatory increases in submaximal pyruvate oxidation. Last, with the exception of IL-6 there were still no increases in TNF-*α* and atrophy markers of cancer cachexia.

## 4. Discussion

Epithelial ovarian cancer is the most lethal gynecological cancer in women. Advanced stages of this disease cause severe muscle weakness, yet the mechanisms remain unknown due in part to the inherent challenges of studying clinical population as well as a limited selection of pre-clinical models. Moreover, it has been suggested that the paucity of metastatic, orthotopic models for cancer cachexia has contributed to failures in clinical trials of therapies designed to preserve muscle mass and/or function that were otherwise based on evidence from other types of preclinical models [15]. Modelling cachexia in a metastatic context is believed to greatly improve the predictive power of preclinical models for identifying mechanisms and therapy development [15]. Here, we developed a new immunocompetent mouse model of metastatic cancer cachexia reflective of late-stage ovarian cancer while retaining the clinically relevant aspects of tumour growth in the nascent organ that can be induced during adulthood. Importantly the findings of this study demonstrate that early muscle weakness precedes clinical signs of ovarian cancer including metastasis and ascites formation as well as atrophy. The eventual development of atrophy seemingly triggers an adaptive response whereby specific force is restored in a sustained manner within limb muscle but only transiently in diaphragm. These pathological and adaptive responses in muscle quality during ovarian cancer coincided with dynamic alterations in pyruvate oxidation and mRNA contents related to numerous mitochondrial pathways but without increases in the cachexia-regulating atrogene programs, TNF-α or GDF15 that have been attributed in this process for other cancers [40,41].

Collectively, the time-dependent responses in force production and fibre size in this new model of ovarian cancer-induced cachexia serve as a foundation for exploring numerous potential mechanisms underlying the development of atrophy-independent and -dependent weakness in relation to metastasis. The findings also highlight how mitohcondrial stress is a defining feature of muscle weakness during ovarian cancer [46].

### 4.1 A novel atrophy-independent weakness in locomotor muscle during ovarian cancer

At the earliest time point assessed (45 Day group), both TA and diaphragm demonstrate lower specific force production. As this measurement is normalized to the size of muscle, any occurrence of atrophy cannot explain this observation. In fact, while modest atrophy was observed in the diaphragm, fibre size was unchanged in the TA at this timepoint. Therefore, the TA demonstrated a pre-atrophy weakness similar to our previous findings in the C26 colorectal cancer model that weakness precedes atrophy in the quadriceps and the diaphragm [16]. Hence, pre-atrophy weakness has now been reported in two distinct models suggesting it is a common phenomenon during cancer. This finding is important because it raises questions regarding the atrophy-independent mechanisms of muscle weakness during cancer – a topic that is considerably understudied in contrast to the literature on atrophy-dependent muscle weakness during cachexia. While it is possible that weakness precedes atrophy in the diaphragm of the current EOC model, future studies would need to examine an earlier time course design.

### 4.2 Reduced mitochondrial pyruvate oxidation is associated with early muscle weakness during ovarian cancer

In the TA, RNAseq identified nuclear genes encoding mitochondrial proteins as the most dominant gene expression stress response early during cancer (45 Day group) when tumours were just appearing, and well before metastasis or atrophy developed. High resolution respirometry revealed that this stress response corresponds with reduced pyruvate oxidation – an index of carbohydrate oxidation – but with no corresponding changes in the capacity for long chain fatty acid oxidation. This finding of a substrate-specific change in respiration was similar in diaphragm at this early time point. There were also no changes in the oxidation of the amino acid-derived substrate glutamate or succinate (generation of FADH_2_ at complex II). As both pyruvate and glutamate generate NADH through their respective dehydrogenases (pyruvate dehydrogenase (PDH) and glutamate dehydrogenase (GDH)), the findings suggest that the ability of Complex I to oxidize NADH may not have been altered in the TA. This suggests that the unique reduction in pyruvate oxidation at 45-days may be due to changes in PDH itself. While there were no changes in PDH or PDH phosphatases mRNA expression identified with RNAseq at this time point (data not shown), future studies could determine if isolated PDH activity is inhibited similar to indications from previous reports in C26 mice [54].

Considering that measurements of oxygen consumption reflect reduction of O_2_ at Complex IV downstream of both Complex I and II, the lack of decreases in both glutamate and succinate oxidation in both TA and diaphragm indicates that the integrated function of the electron transport chain was not altered, at least as could be detected within the physiologically relevant context of ADP-stimulated (coupled) respiration. This finding is interesting given that protein contents of specific subunits of ETC complexes were not changed across time points. In the 75 Day and 90 Day groups, the protein contents of ETC protein subunits measured by western blot were not changed, but mRNA content of these subunits were decreased (data not shown; C1 - NDUFB8, CII – SDHB, CIII – UQCRC2, CV – ATP5A). While speculative, these findings suggest a number of possibilities including increased ETC protein stability or that reductions in mRNA reflect increased translation rather than decreased gene expression [55].

Given that ovarian cancer does not reduce oxidation of the other substrates explored at this early timepoint, it is also difficult to define the unique reductions in pyruvate oxidation as a ‘mitochondrial dysfunction’ *per se*. This is a critical outcome of the present investigation and highlights the importance of comparing substrates to each other and to defined primary outcomes of myopathy tracked over time, and across muscle types. This approach determines whether mitochondrial stress responses affects a central governance of oxidative phosphorylation or an adaptive reprogramming - a concept and perspective we have proposed previously for the study of myopathies [56].

### 4.3 Recovery of specific force during the development of atrophy is more sustained in limb muscle versus diaphragm

As atrophy developed by the 75 Day group in both muscles, specific force partially recovered despite the appearance of atrophy in the TA with even greater atrophy in the diaphragm. As specific force is a measure that is independent of muscle mass, this finding raises questions regarding the mechanism of intrinsic improvements within the atrophied muscle itself. This remarkable adaptive response in both muscles diverged in the 90 Day group. Particularly, specific force recovered even further in the TA, whereas diaphragm force plummeted to very low levels.

Unlike the 45 Day group, changes in force production were not consistently related to changes in pyruvate oxidation given this function remained low as force recovered at 75 days, with increases in pyruvate oxidation at 90 days being positively or inversely related to muscle force production in the TA and diaphragm, respectively. Nonetheless, decrements in pyruvate oxidation were more strongly related to pre-atrophy weakness early during cancer. This finding warrants further examination with targeted approaches that determine whether altered glucose metabolism contributes to weakness uniquely at early stages of ovarian cancer.

As the mechanisms underlying the progressive vs transient increase in force production in both muscles require further investigation, RNAseq analyses in the TA demonstrated mRNA contents related to chromatin regulation and biosynthetic processes that can be explored for the generation of numerous hypotheses in future investigations. Likewise, increased mRNA contents corresponding to genes related to myofibril assembly and actomyosin structure organization during the pre-atrophy period at 45 days in the TA suggests potential turnover of sarcomeric structures may have occurred. Future studies could consider whether pre-atrophy weakness was due to declining quality of contractile machinery. Last, Reactome enrichment analysis identified 19 “muscle contraction” genes upregulated at the 90-day time point in the TA. Some of these genes are related to calcium handling (ATP2b4, ITPR2, RyR1, and ATP2a1), suggesting increases in force production could be related to calcium regulation. While RNAseq was not performed in the diaphragm, Rt-PCR analysis did identify decreases in RyR and SERCA mRNA content concurrent with a decrease in force-frequency production. Functional calcium handling measures could be performed in the future to explore potential relationships between mitochondrial ATP supply supported by carbohydrate oxidation and the energetic cost of contraction [57].

### 4.4 Mitochondrial-cytoplasmic phosphate shuttling: two systems, two different responses during ovarian cancer

This study was also designed to consider how mitochondria shuttle phosphate to the cytoplasm in the form of both PCr (Phosphocreatine) and ATP in order to gain deeper insight into the precise mechanisms by which skeletal muscle mitochondria demonstrate metabolic reprogramming (as explained in *Results* and in [46]). These modeling approaches identify early reductions in pyruvate oxidation in both TA and diaphragm that were observed more consistently in the dominant creatine-dependent pathway (PCr export) yet both phosphate shuttling systems (ATP and PCr export) improved over time. The late stage increases in both systems, with an apparent supercompensation in the creatine-independent (ATP) shuttling system, indicate a mitochondrial ‘hormesis’ consistent with a perspective that mitochondria attempt to enhance the supply of ATP to a failing muscle fibre as the stress of cancer intensifies.

Collectively, this experimental design led to findings that can guide pre-clinical therapy development to treat cancer-induced muscle weakness. For example, the relationships would support further investigation into therapies that preserve pyruvate oxidation or creatine-dependent metabolism early in cancer could be explored to determine if the pre-atrophy weakness can be prevented.

### 4.5 mH_2_O_2_ emission: a delayed relationship with weakness?

mH_2_O_2_ emission was not elevated at the early 45 Day timepoint corresponding to muscle weakness in both muscles. Therefore, there was a stronger relationship between early weakness and reduced pyruvate oxidation and creatine-dependent respiration in both muscles than to oxidative stress. Rather, mH_2_O_2_ emission was increased by the 75 Day group in both muscles. By the 90 Day group, mH_2_O_2_ emission returned to control levels in both muscles although this depended on the pathway assessed. While mH_2_O_2_ emission derived from the reverse electron transfer pathway was more consistently elevated in both muscles, the unique time-dependent patterns of both systems further highlight the complexities of mitochondrial reprogramming that would not be captured by traditional single pathway analyses. Collectively, there is a clear increase in mH_2_O_2_ in mid to late stages of cancer corresponding to atrophy. Therefore, the findings serve as a basis for examining the potential roles of mitochondrial-derived redox signals in regulating muscle fibre size distinct from mechanisms governing earlier muscle weakness that was more strongly related to changes in oxidative phosphorylation in this model. The findings also suggest that mitochondrial-targeted antioxidants could be tested to determine if these mH_2_O_2_ responses are partially contributing to atrophy during ovarian cancer. Indeed, a previous study in C26 cancer mice demonstrated the mitochondrial cardiolipin-targeting small peptide SS-31 prevented atrophy at later stages of development in relation to lower mH_2_O_2_ emission [58].

### 4.6 Mechanisms regulating cachexia in an orthotopic, metastatic, epithelial ovarian cancer model appear to differ from other pre-clinical models

Contemporary theories proposes muscle wasting during cancer cachexia is induced by circulating factors generated by the host or tumour which trigger protein degradation and loss of myofibrillar proteins [4,59,60]. Several genes and cytokines are thought to regulate this skeletal muscle degradation but atrogin, Murf-1, TNF-a, IL-6, and GDF15 are perhaps most commonly identified and measured [40,41]. However, the current investigation using an immunocompetent, orthotopic, metastatic model of ovarian cancer does not demonstrate robust activation of these pathways. Indeed, with the exception of IL-6 in the diaphragm, the factors regulating muscle atrophy seem to be different within the current model. The time-specific increases in IL-6 in the diaphragm could be explored given this cytokine is integral for the development of muscle loss in the *Apc^min/+^* mouse (genetic spontaneous colorectal cancer model) [61,62].

The absence of atrogene responses is similar to a patient-derived xenograft (PDX) model whereby, pancreatic ductal adenocarcinoma (PDAC) tumours from cancer patients are orthotopically injected into immunodeficient NSG mice [63]. Within this model, the TA demonstrates up-regulation in canonical atrophy-associated pathways (ubiquitin-mediated protein degradation), while the diaphragm demonstrates an up-regulation in genes related to the inflammatory response [63]. This could suggest that orthotopic models demonstrate distinct cachexia profiles between muscles that are unique to ectopic models.

Reductions in food intake and physical activity are thought to contribute partially to cachexia [64,65]. The degree to which these patterns contribute to pre-atrophy weakness or atrophy itself in the current study requires further investigation. However, previous work in C26 mice has shown that reductions in food intake did not contribute to reduced muscle weights, fiber CSA, or muscle force given pair-fed mice retained normal muscle parameters compared to tumour bearing mice (35, 36).

### 4.7 Perspectives and Conclusions

The discovery that muscle weakness and mitochondrial stress precedes metastasis and ascites accumulation in ovarian cancer raises the question of whether pre-atrophy weakness could be an early diagnostic marker of cancer, given that ovarian malignancy is largely undetectable at premetastatic stages. Indeed, most women are diagnosed with ovarian cancer at stage III – a time where metastasis/ascites have already started, and survival rates are low [17,18,66]. Thus, early cancer detection is suggested to be one of the best strategies for cancer prevention [67]. Exploring this possibility would be complex given muscle weakness and altered mitochondrial functions could occur in other health conditions such as ageing and muscle disuse [68,69]. These findings also position mitochondrial reprogramming as a potential therapeutic target in pre-atrophy weakness and cachexia during metastatic ovarian cancer.

In conclusion, this is the first mouse model of epithelial ovarian cancer-induced muscle weakness that offers the advantages of orthotopic injections of EOC cells into the ovarian bursa that can be performed in immunocompetent mice during adulthood. The model also demonstrates the critical clinical feature of metastasis in the abdominal cavity similar to what occurs in women with late-stage ovarian cancer. The identification of an early muscle weakness that precedes both atrophy and metastasis provide a new direction for research in understanding the atrophy-independent mechanisms of muscle weakness during ovarian cancer. We identified substrate-specific alterations in mitochondrial oxidative phosphorylation and increases in mitochondrial reactive oxygen species that coincide with early pre-metastatic weakness. The model also demonstrates late-stage compensatory relationships between mitochondrial metabolism and specific force restoration in limb muscle suggesting a remarkable adaptive mechanism that appears to be muscle-specific. The time-dependent and muscle-specific relationships described in this new model provide will support continued efforts in defining atrophy-independent and -dependent mechanisms of weakness during ovarian cancer in relation to metastasis and for guiding the design of pre-clinical therapy development investigations.

## List of Abbreviations

-Creatine: Without creatine
+Creatine: With creatine
ADP: Adenosine diphosphate
ATP: Adenosine triphosphate
BLEB: Blebbistatin
BW: Body weight
C26: Colon-26
Cr: Creatine
CSA: Cross sectional area
CTRL: Control
DEG: Differentially expressed gene
eMHC: Embryonic myosin heavy chain
EOC: Epithelial ovarian cancer
ETC: Electron transport chain Extensor digitorum longus (EDL)
FADH_2_: Flavin adenine dinucleotide
GA: Gastrocnemius
GDF15: Growth differentiation factor 15
GDH: Glutamate dehydrogenase
GO: Gene ontology
IL-6: Interleukin 6
L_o_: Optimal Resting Length
mH_2_O_2_: Mitochondrial hydrogen peroxide
MHC: Myosin Heavy Chain
mtCK: Mitochondrial creatine kinase
MURF-1: Muscle RING-finger protein-1
NADH: Nicotinamide adenine dinucleotide
OCT: Optimal cutting temperature
OXPHOS: Oxidative phosphorylation
PBS: Phosphate buffered saline
PCr: Phosphocreatine
PDAC: Pancreatic ductal adenocarcinoma
PDH: Pyruvate dehydrogenase
PLT: Plantaris
PmFB: Permeabilized Fiber bundles
Quad: Quadriceps
RET: Reverse electron transfer
RNASeq: RNA sequencing
Rt-PCR: Reverse transcription-polymerase chain reaction
RyR1: Ryanodine receptor
SERCA: Sarcoendoplasmic reticulum calcium ATPase
Sol: Soleus
TA: Tibialis anterior
TNFα: Tumour necrosis factor alpha

## Authors’ Contributions

**Luca J. Delfinis:** conceptualization, methodology, validation, formal analysis, investigation, writing -original draft, writing – review & editing, visualization, project administration; **Leslie M. Ogilvie:** conceptualization, methodology, investigation, writing – review & editing; **Shahrzad Khajehzadehshoushtar**: conceptualization, methodology, investigation, writing – review & editing; **Shivam Gandhi:** investigation and writing – review & editing; **Madison C. Garibotti:** investigation and writing – review & editing; **Arshdeep K. Thuhan**: investigation and writing – review & editing; **Kathy Matuszewska:** methodology; **Madison Periera**: methodology; **Ronald G. Jones III:** formal analysis, data curation and writing – review & editing; **Arthur J. Cheng**: validation and writing – review & editing; **Thomas J. Hawke**: validation and writing – review & editing; **Nicholas P. Greene**: validation and writing – review & editing; **Kevin A. Murach**: formal analysis, validation and writing – review & editing; **Jeremy A. Simpson:** conceptualization and writing – review & editing; **Jim Petrik:** conceptualization, methodology and writing – review & editing; **Christopher G.R. Perry:** conceptualization, methodology, validation, writing -original draft, writing – review & editing, visualization, project administration, supervision, funding acquisition.

## Acknowledgment

Funding was provided to CGRP by the National Science and Engineering Research Council (NSERC, 436138-2013 and 2019-06687) and an Ontario Early Research Award (2017-0351) with infrastructure supported by the Canada Foundation for Innovation, the Ontario Research Fund, and the James H. Cummings Foundation. LJD and LMO were supported by NSERC CGS-D scholarship. SK was supported by NSERC CGS-M scholarship. SG and AKT were supported by Ontario Graduate Scholarship (OGS). KAM was supported by National Institute of Health (NIH) R01 AG080047. JP was supported by Canadian Institute of Health Research (CIHR) 450209.

## Supplemental Figure Legends

**SFigure 1.**
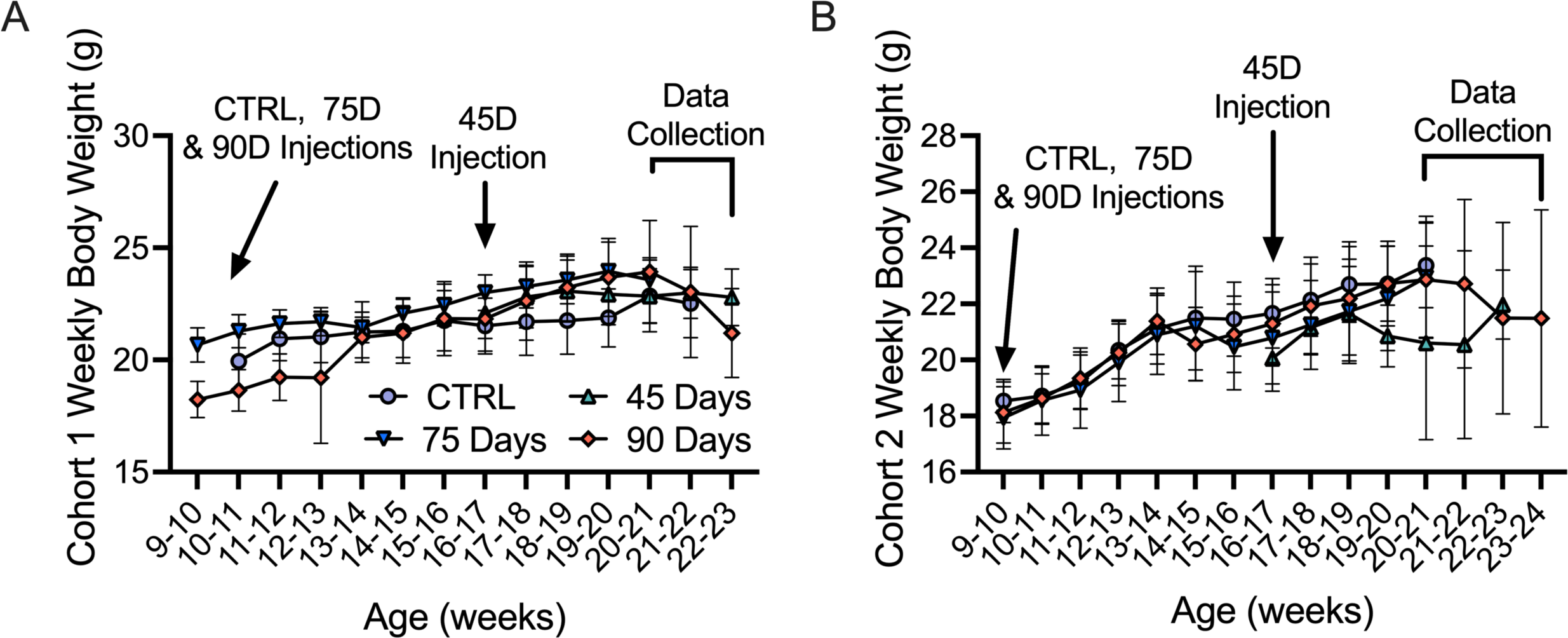
Weekly body weights and age of EOC injections throughout study. This study used two mice per “n” from separate cohorts to obtain enough tissue to complete all experiments. Weekly body weights were measured in mice from cohort 1 (A, n =12) and cohort 2 (B, n =12). Results represent mean ± SD. All data was analyzed using a one-way ANOVA and followed by a two-stage step-up method of Benjamini, Krieger and Yukutieli multiple comparisons test. C57BL/6J female mice ∼75 days post PBS injection as controls (CTRL); C57BL/6J female mice ∼45 days post ovarian cancer injection (45 Days); C57BL/6J female mice ∼75 days post ovarian cancer injection (75 Days); C57BL/6J female mice ∼90 days post ovarian cancer injection (90 Days).

**SFigure 2.**
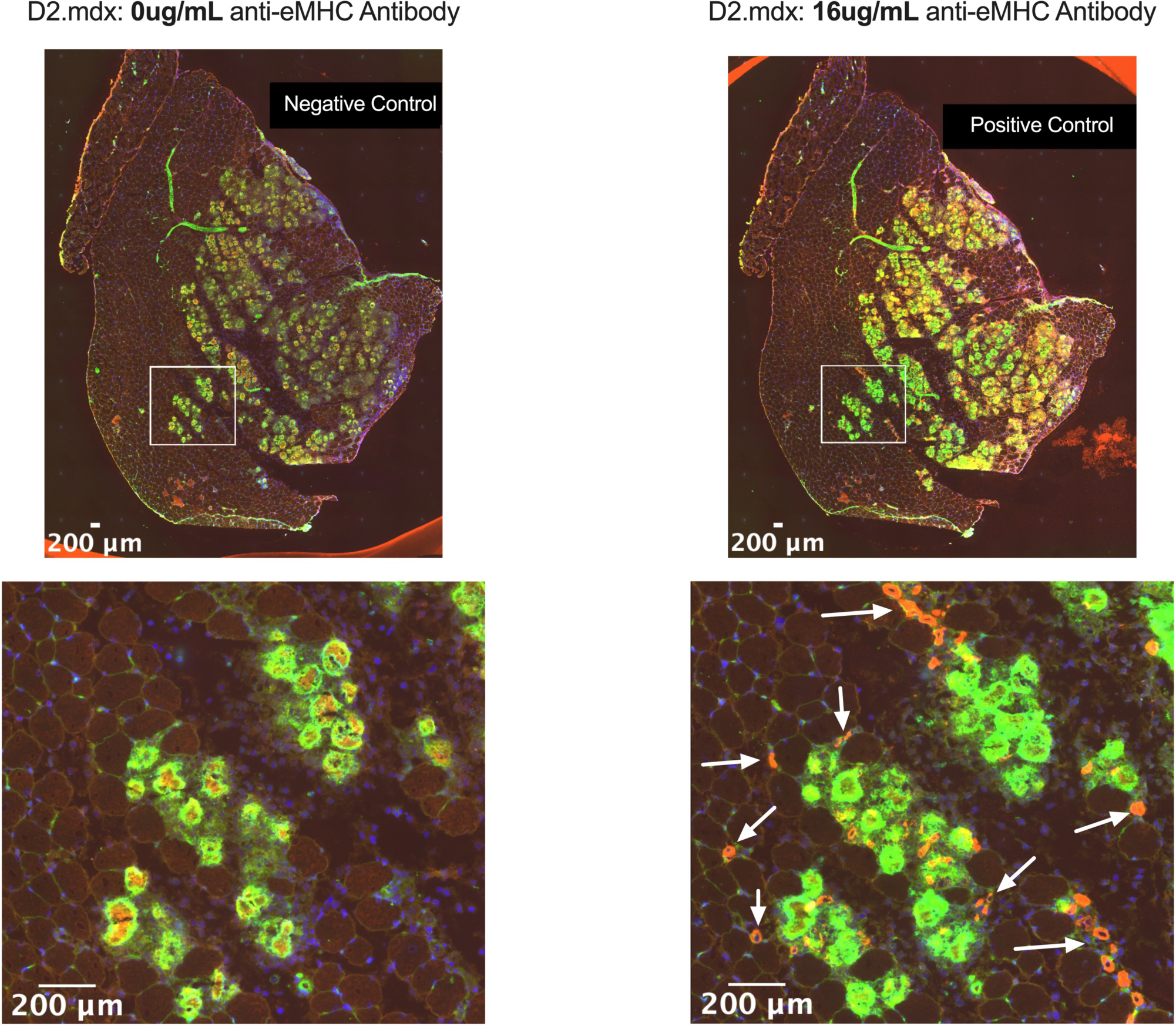
Positive and negative control experiments of eMHC protocol. Tibialis anterior muscle from D2.mdx mice were used as a positive control to validate the eMHC histology technique. Technical replicates of the same tissue were incubated with no eMHC antibody (left) and with 16*μ*g/mL of eMHC primary antibody (right).

**SFigure 3.**
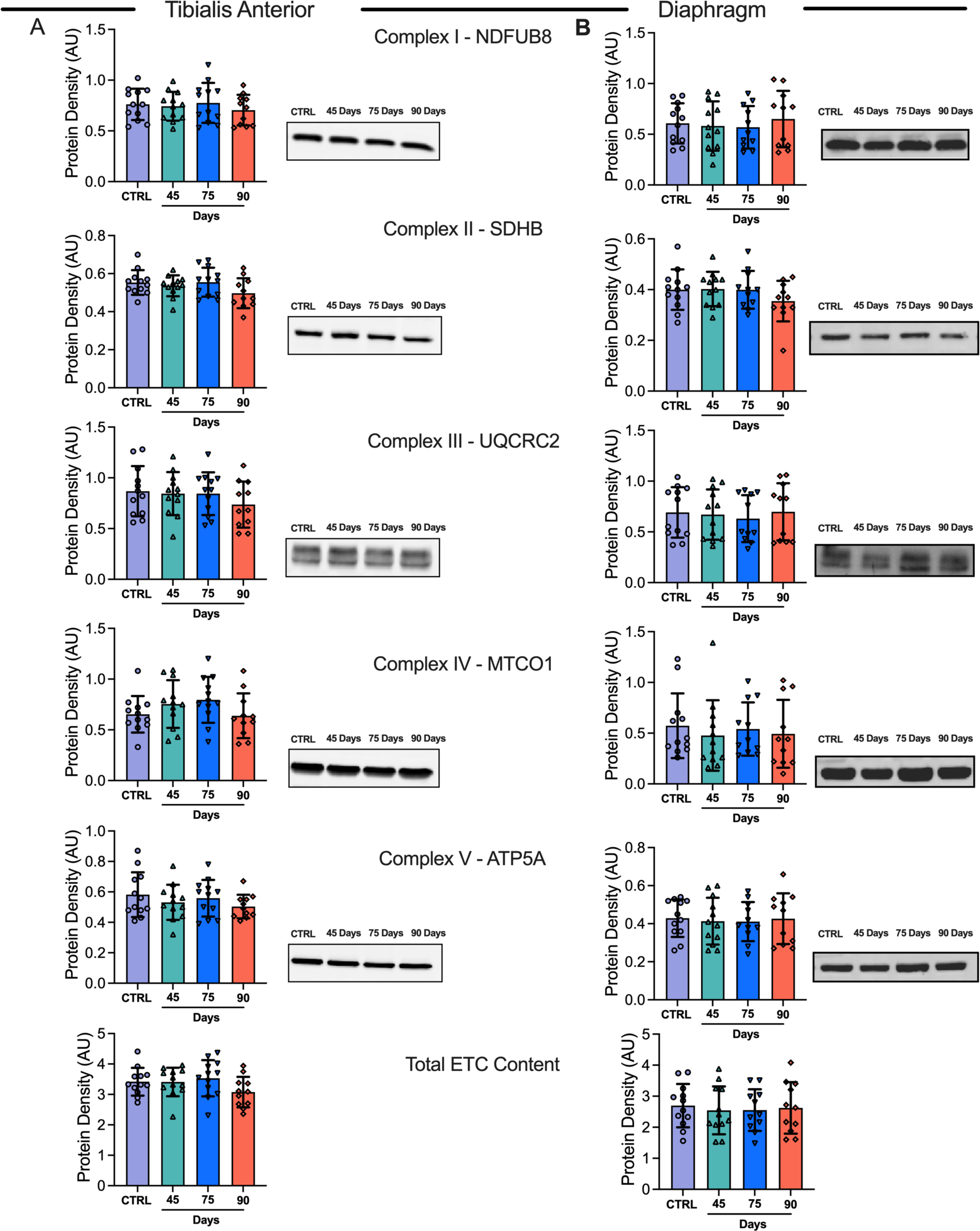
Muscle-specific evaluation of electron transport chain (ETC) complex subunit markers in EOC injected tibialis anterior and diaphragm skeletal muscle. Protein content of ETC subunits was quantified in the tibialis anterior (A, n = 12) and diaphragm (B, n = 12) Results represent mean ± SD. All data was analyzed using a one-way ANOVA or Kruskal-Wallis test when data did not fit normality. All ANOVAs were followed by a two-stage step-up method of Benjamini, Krieger and Yukutieli multiple comparisons test. C57BL/6J female mice ∼75 days post PBS injection as controls (CTRL); C57BL/6J female mice ∼45 days post ovarian cancer injection (45 Days); C57BL/6J female mice ∼75 days post ovarian cancer injection (75 Days); C57BL/6J female mice ∼90 days post ovarian cancer injection (90 Days).

**SFigure 4.**
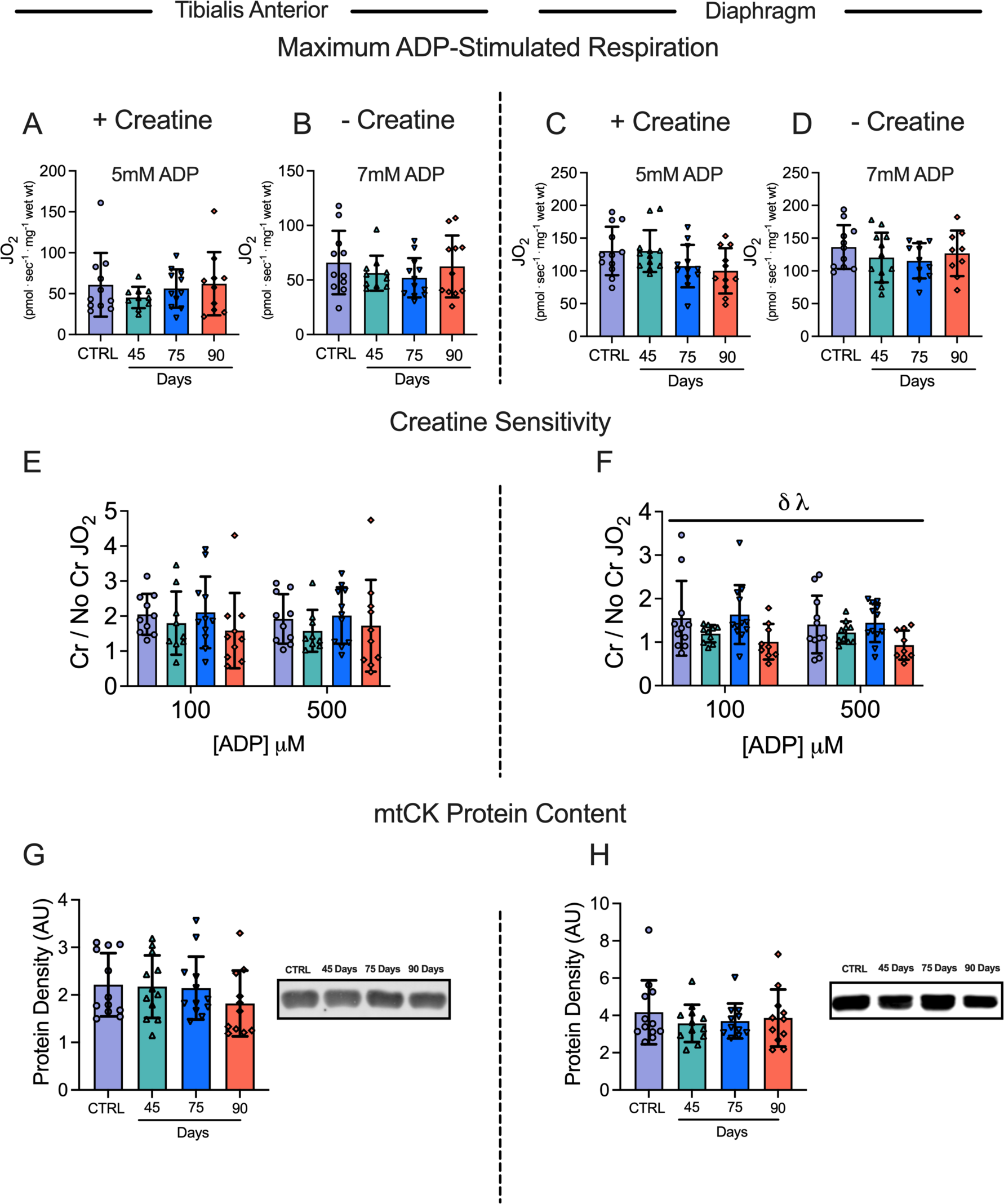
**Maximum ADP-stimulated respiration, creatine sensitivity ratios and mitochondrial creatine kinase (mtCK) protein content in tibialis anterior and diaphragm muscle of EOC injected mice**. Maximum ADP-stimulated mitochondrial respiration was evaluated in the tibialis anterior and diaphragm both in the presence and absence of creatine (A-D, n = 9-12). A ratio of +Creatine/-Creatine respiration in the tibialis anterior and diaphragm muscle was generated at 100μM and 500μM (apparent Km of mtCK) as an index of creatine sensitivity (E & F, n = 9-12). mtCK protein content was also quantified in both muscles (n = 12). Results represent mean ± SD. λ *p* < 0.05 75 Day vs 90 Day; δ *p* < 0.05 Control versus 90 Day. Figures A-D, G and H were analyzed using a one-way ANOVA or Kruskal-Wallis test when data did not fit normality. Figures E and H were analyzed using a two-way ANOVA (main effect shown only). All ANOVAs were followed by a two-stage step-up method of Benjamini, Krieger and Yukutieli multiple comparisons test. C57BL/6J female mice ∼75 days post PBS injection as controls (CTRL); C57BL/6J female mice ∼45 days post ovarian cancer injection (45 Days); C57BL/6J female mice ∼75 days post ovarian cancer injection (75 Days); C57BL/6J female mice ∼90 days post ovarian cancer injection (90 Days).

**SFigure 5.**
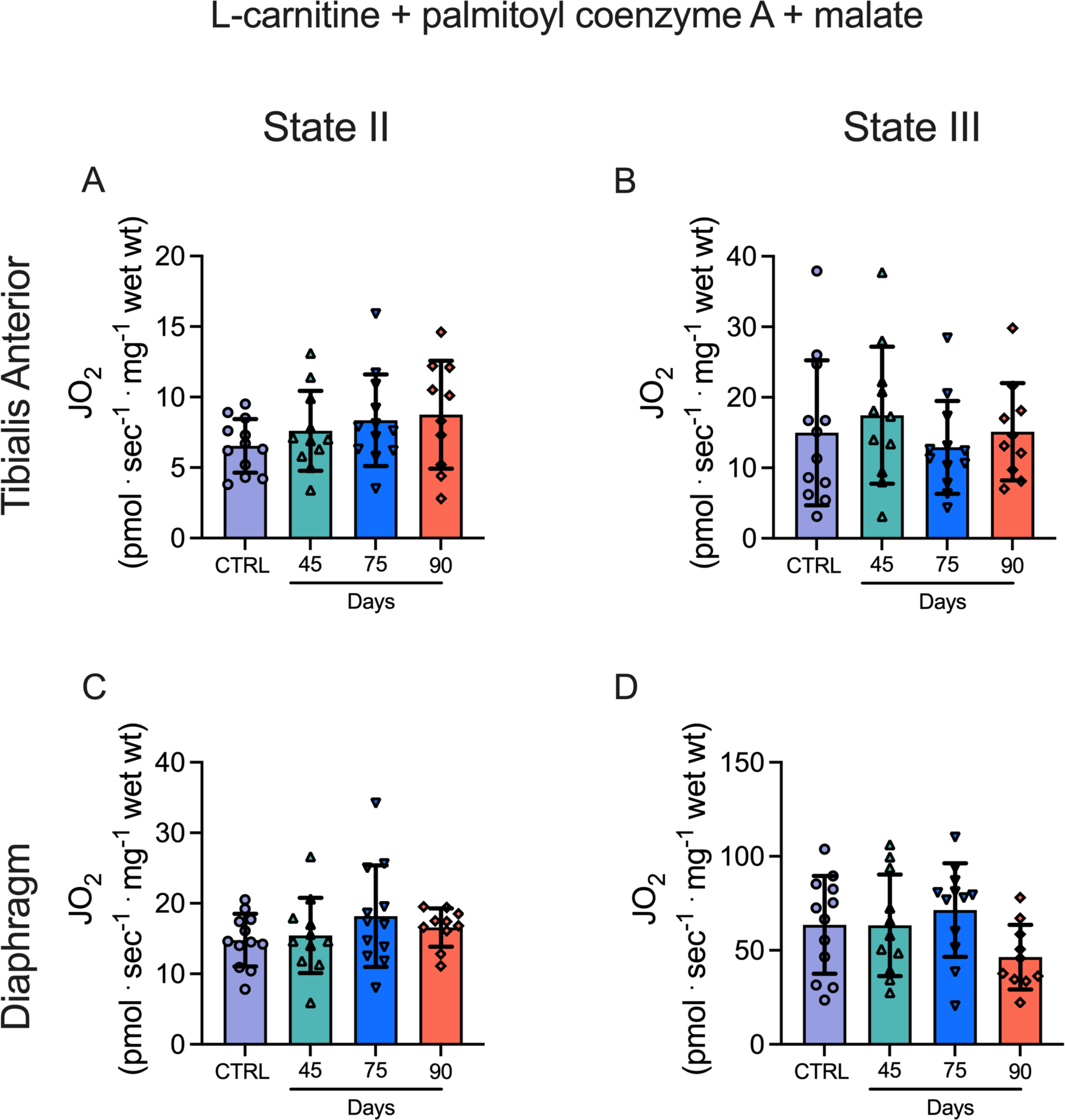
Fatty acid-supported mitochondrial respiration in tibialis anterior and diaphragm of EOC injected mice. State II (L-carnitine + palmitoyl coenzyme A + malate; absence of ADP) mitochondrial respiration was evaluated in the tibialis anterior and diaphragm muscle in the presence of 20mM creatine (A & C, n = 10-12). State III (5mM ADP) mitochondrial respiration was also evaluated in TA and diaphragm muscle (B & D, n =10-12) Results represent mean ± SD. All data was analyzed using a one-way ANOVA or Kruskal-Wallis test when data did not fit normality. All ANOVAS were followed by a two-stage step-up method of Benjamini, Krieger and Yukutieli multiple comparisons test. C57BL/6J female mice ∼75 days post PBS injection as controls (CTRL); C57BL/6J female mice ∼45 days post ovarian cancer injection (45 Days); C57BL/6J female mice ∼75 days post ovarian cancer injection (75 Days); C57BL/6J female mice ∼90 days post ovarian cancer injection (90 Days).

**SFigure 6.**
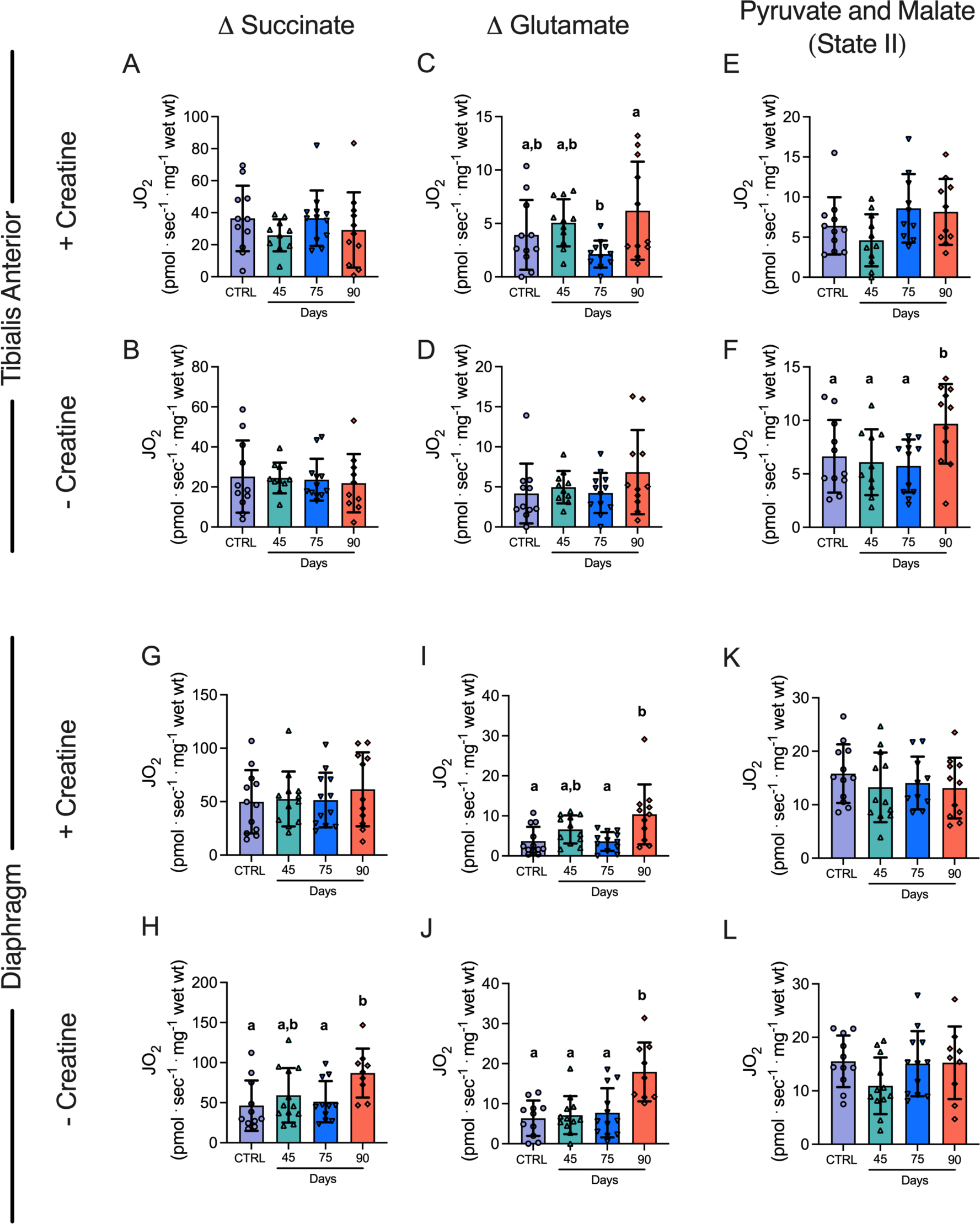
Multiple substrate evaluation of oxygen consumption in tibialis anterior and diaphragm of EOC injected mice. Oxygen consumption was evaluated in tibialis anterior bundles using succinate both in the presence and absence of creatine (A & B). Glutamate-supported respiration was also evaluated in the presence and absence of creatine (C & D). State. II (absence of ADP) was also evaluated in the presence and absence of creatine (E & F). This was repeated in the diaphragm (G-L). Results represent mean ± SD. n = 9-12. Lettering denotes statical significance when different from each other (*p* < 0.05). All data was analyzed using a one-way ANOVA or Kruskal-Wallis test when data did not fit normality. All ANOVAs were followed by a two-stage step-up method of Benjamini, Krieger and Yukutieli multiple comparisons test. C57BL/6J female mice ∼75 days post PBS injection as controls (CTRL); C57BL/6J female mice ∼45 days post ovarian cancer injection (45 Days); C57BL/6J female mice ∼75 days post ovarian cancer injection (75 Days); C57BL/6J female mice ∼90 days post ovarian cancer injection (90 Days).

**SFigure 7.**
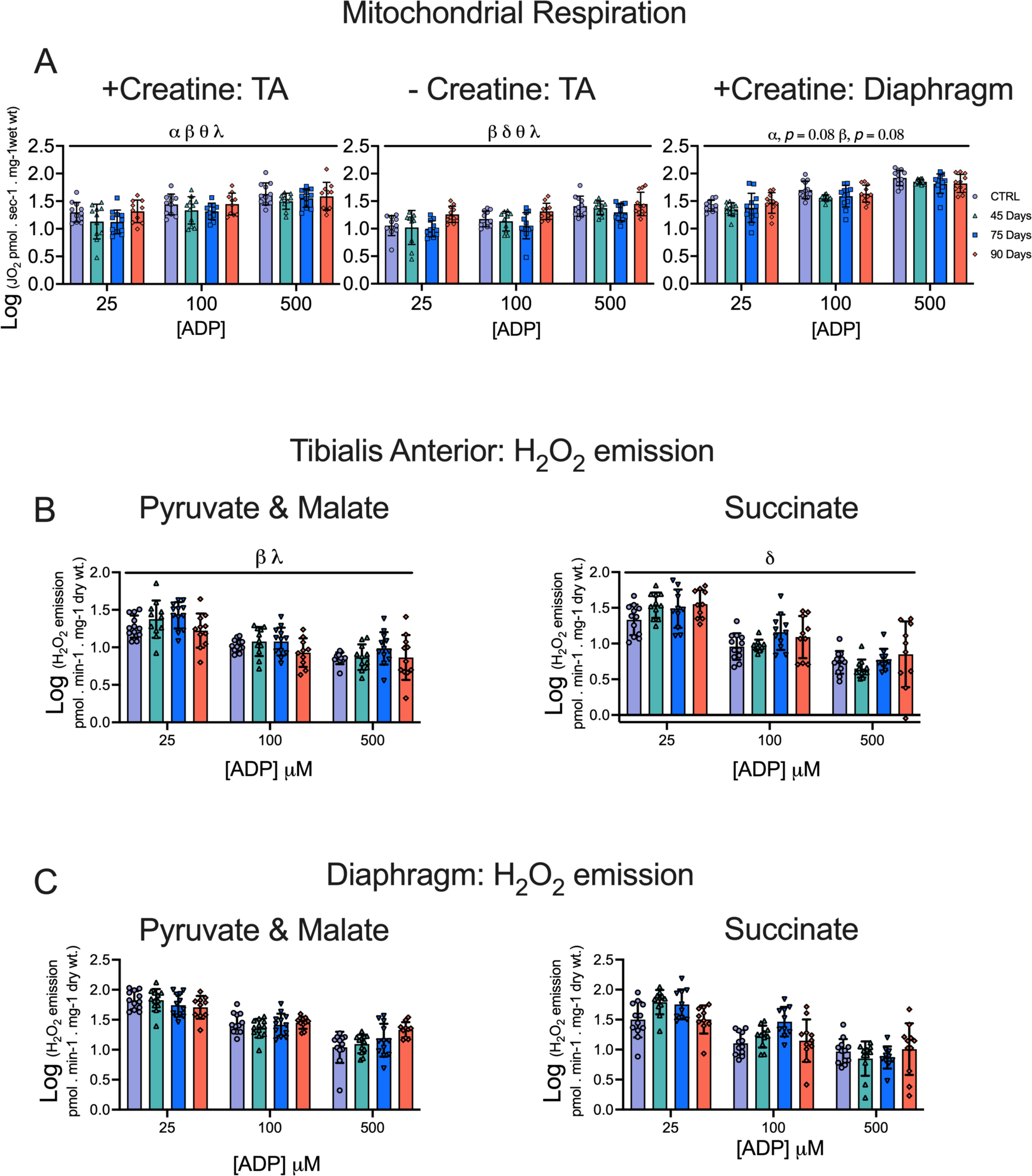
Log transformed data for analysis in tibialis anterior and diaphragm that did not fit a normal distribution. Data that did not fit normality were log transformed and then analyzed using standard 2-way ANOVAs. Results represent mean ± SD. n = 9-12. α *p* < 0.05 Control versus 45 Day; β *p* < 0.05 Control versus 75 Day; δ *p* < 0.05 Control versus 90 Day; θ *p* < 0.05 45 Day versus 90 Day; λ *p* < 0.05 75 Day vs 90 Day. All Data were analyzed using a two-way ANOVA. All ANOVAs were followed by a two-stage step-up method of Benjamini, Krieger and Yukutieli multiple comparisons test. C57BL/6J female mice ∼75 days post PBS injection as controls (CTRL); C57BL/6J female mice ∼45 days post ovarian cancer injection (45 Days); C57BL/6J female mice ∼75 days post ovarian cancer injection (75 Days); C57BL/6J female mice ∼90 days post ovarian cancer injection (90 Days).

**STable 1.**
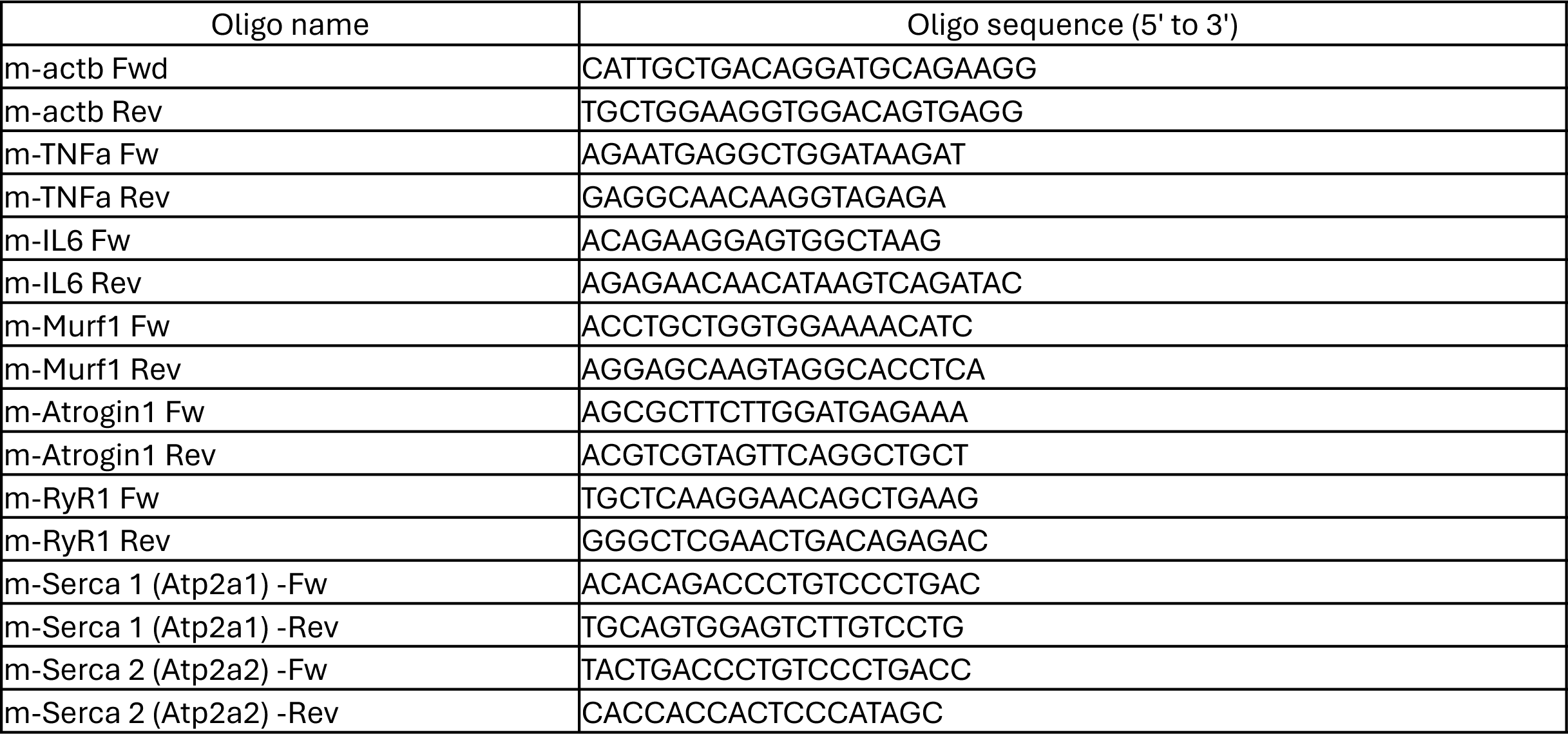
List of primers used for qtPCR.

